# Structural and mechanistic analyses reveal collaborative regulation of HSF1 by the Hsp70-Hsp90 chaperone systems

**DOI:** 10.64898/2026.08.27.747592

**Authors:** Tristan W. Owens, Kaitlin Schaefer, Trenton M. Peters-Clarke, James A. Wells, David A. Agard

## Abstract

Heat shock factor 1 (HSF1) is the master transcriptional regulator of cellular response to disrupted cytosolic protein homeostasis. Temperature change, oxidation, and other stresses drive the trimerization and activation of HSF1 to induce expression of molecular chaperones, such as heat shock proteins Hsp70 and Hsp90, which sit at the center of cellular proteostatic networks. In turn, the HSPs and co-chaperones regulate HSF1, but mechanistic details of this cycle remain largely unknown. We developed a FRET-based approach to simultaneously monitor HSF1 conformational change and oligomeric state throughout activation and inactivation. By reconstituting Hsp-HSF1 interactions *in vitro*, we find that monomerization of HSF1 resembles fibril disassembly through coordinated Hsp40-Hsp70 activity. We then used site-specific photocrosslinking to track HSF1 loading into Hsp90 complexes, Hsp90 cycling, and stress-induced shifts in Hsp90-HSF1 interactions. Whereas Hsp90 inhibitors force ‘loading state’ type Hsp90-HSF1 interactions, heat shock promotes faster Hsp90 cycling. In this reconstituted system, HSF1-Hsp90 interactions are unexpectedly strongly dependent on the co-chaperone HOP, in contrast to canonical Hsp90 clients. Combining cryo-EM structures of Hsp90-HSF1 loading state and maturation state complexes with crosslinking mass spectroscopy and biophysical experiments, we show that Hsp90 holds HSF1 in a pre-activated, extended monomer state that is primed for trimerization. We propose this state both enhances responsivity but also promotes cytoplasmic-nuclear shuttling through exposure of the NLS. Notably, Hsp90-bound HSF1 can trimerize and bind DNA, placing it on-pathway for transcriptional activation. Together, our results unify previously contradictory view on Hsp90’s role in HSF1 regulation. The integrated combination of in vitro reconstitution, photocrosslinking, cryoEM and MS is an exciting new paradigm for the study of many dynamic systems including other complex proteostasis components.

## Introduction

The cellular machinery to maintain protein homeostasis is conserved across all domains of life and enables organisms to adapt to environmental and cellular stresses. A central aspect of this response is induction of heat shock proteins (HSPs), such as Hsp40, Hsp70, and Hsp90, that act as molecular chaperones to facilitate protein folding, protect against aggregation, and aid the degradation of misfolded proteins (Vabulas, 2010)^1^. In eukaryotes, induction of HSPs in response to cytosolic stress is coordinated by the master regulator heat shock transcription factor 1 (HSF1) (Gomez-Pastor, 2018)^2^. The transcriptional program controlled by HSF1, the heat shock response (HSR), entails the upregulation of hundreds of genes and downregulation of thousands of others (Mahat, 2016; Himanen, 2022)^3,4^. Broadly, HSF1 activity is thought to be controlled in turn by HSPs and their co-chaperones, providing a feedback mechanism that establishes a sufficiency of proteostatic capacity.

HSF1 and the HSPs also play critical roles in unstressed cells, and basal expression of HSPs is also typically dependent on HSF1 (Solís 2014; Takii 2024)^5,6^. Concordantly, reduced HSF1 activity is common in neurodegenerative diseases characterized by protein misfolding (Neef, 2011; Rozema 2025)^7,8^. Through mechanisms that are independent of the HSR, but less well studied, HSF1 signaling also promotes cell growth, proliferation, and metabolism (Barna, 2019)^9^. At the same time Hsp90, facilitates the folding and activation of ∼10% of the proteome, including 60% of human kinases and steroid hormone receptors (Schopf, 2017)^10^. Tumor cells frequently depend on heightened levels of HSPs for both the chaperoning of oncoproteins and maintaining proteostasis during rapid growth and metabolic imbalance (Jaeger, 2019)^11^. Thus, HSF1 is critical to tumor development and progression, and HSF1 activity is strongly correlated with poor prognosis in common cancers (Santaga, 2011; Mendillo, 2012; Dai, 2016)^12–14^. Many efforts have been made to develop cancer therapies targeting Hsp90, but a key confounding factor is the interlinked regulation of HSF115. ATP-competitive Hsp90 inhibitors induce the HSR, upregulating HSPs which can support the cancer. Resolving the details of HSF1 activation and regulation will help us understand its function in both health and disease.

Heat shock and other proteotoxic stresses convert inactive, monomeric HSF1 to an active, trimeric state that translocates to the nucleus and binds heat shock elements (HSEs) in the promoters of target genes (Figure 1A) (Gomez-Pastor, 2018)^2^. HSF1 comprises an amino-terminal DNA binding domain (DBD) (Vuister, 1994; Harrison, 1994; Neudegger, 2016)^16–18^, a leucine zipper domain (Lz1-3) (Sorger, 1989; Clos, 1990; Rabindran, 1993)^19–21^, an unstructured regulatory domain (RD) (Green, 1995)^22^, a second leucine zipper domain (Lz4), and a unstructured carboxy-terminal transactivation domain (TAD) (Green, 1995)^22^. In the resting state, HSF1 is autoinhibited by interactions between the Lz1-3 and Lz4 domains; during activation Lz4 is displaced and HSF1 trimerizes by formation of an intermolecular coiled-coil between Lz1-3 domains (Figure 1B) (Hentze, 2016)^23^. Trimeric HSF1 has a much higher binding affinity for tripartite HSEs, and it then recruits additional transcription factors and RNA polymerase via the TADs.

**Figure 1:**
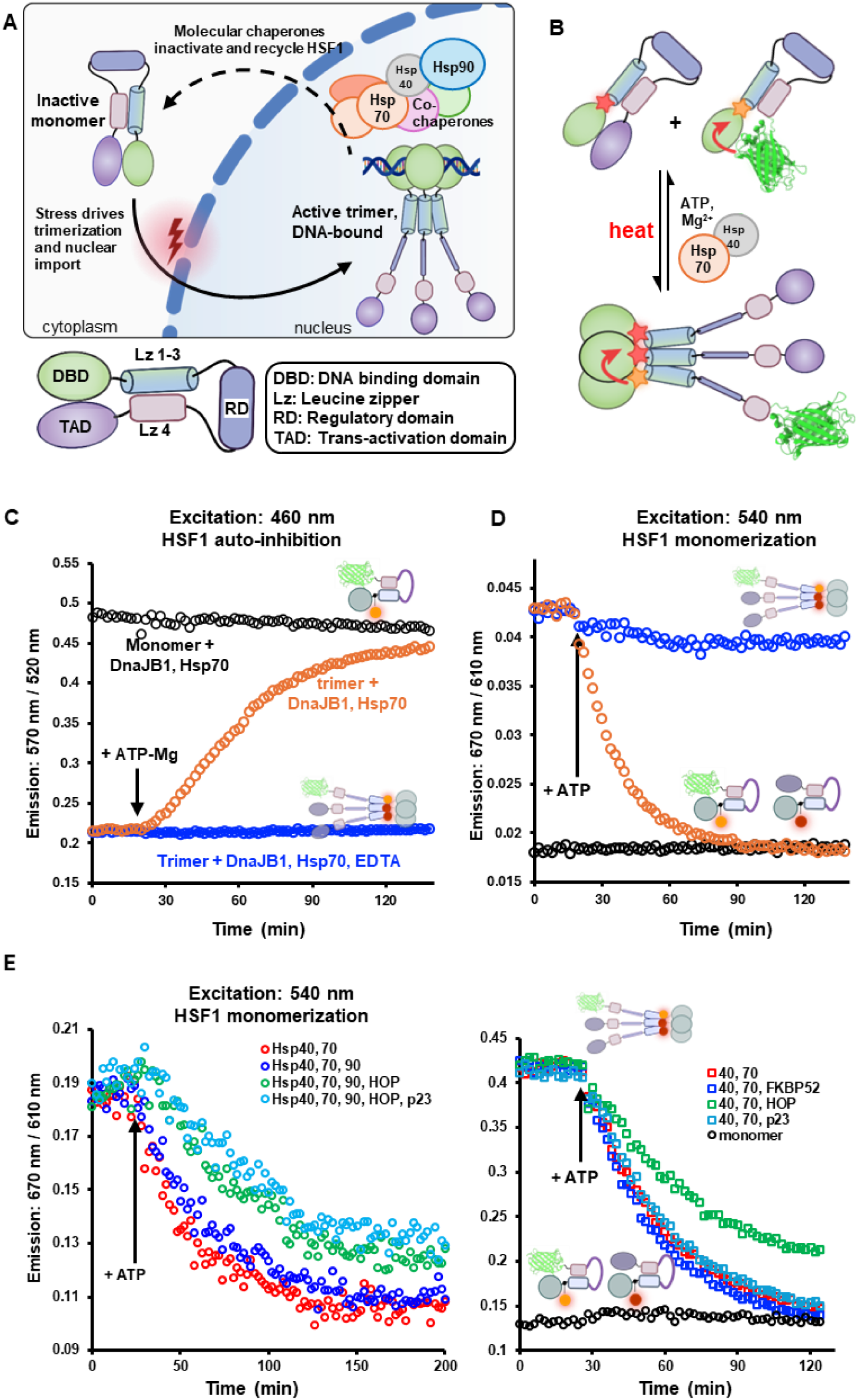
Using a FRET-based approach to probe HSF1 activation and chaperone-mediated regulation. (A) In a simple schematic, HSF1 cycles between inactive, cytoplasmic monomers and nuclear, DNA bound trimers. This equilibrium is regulated by stress and chaperones. Below, the five domains of HSF1. (B) A biochemical system with two HSF1 constructs was designed for FRET detection of autoinhibition (GFP to Alexa546, orange star), and oligomerization (Alexa546 to Alexa647, red star). (C) Monitoring HSF1auto-inhibition shows that DnaJB1 (3 µM), Hsp70 (8 µM), and ATP-Mg restore HSF1 trimers to an auto-inhibited state, not release autoinhibition of HSF1 monomers. (D) HSF1 oligomerization signal shows that DnaJB1, Hsp70 converts HSF1 trimers to monomers in an ATP-dependent manner. (E) Hsp90 (12 µM) + HOP (6 µM) or HOP alone slows HSF1 monomerization by Hsp40 and Hsp70.

Previous work implicated Hsp70, Hsp90, and co-chaperones in both the activation and inactivation of HSF1. Until recently, the dominant model for HSF1 regulation was the chaperone titration model, that a multichaperone complex maintained inactive HSF1, and that protein folding stress competed these chaperones away from HSF1 allowing its activation (Anckar, 2011)^24^. A simple model is supported in yeast (Masser, 2019)^25^, but the human system is more complicated. In a biochemical reconstitution, Kmiecik and co-workers (2020)^6^ demonstrated DnaJB1 and Hsp70 are sufficient to disassemble human HSF1 trimers but did not see a role for Hsp90. Variously, Hsp90 has been reported to repress activation of HSF1 monomers (Zou, 1998)^27^, to inactivate HSF1 trimers by removing them from DNA (Ali, 1998; Kijima, 2018)^28,29^ or blocking transcription, and to promote trimerization of monomers (Hentze, 2016)^23^. Nonetheless, Hsp90-HFS1 complexes have proved difficult to isolate, typically requiring chemical crosslinking or inactivating mutations in Hsp90. That Hsp90 inhibitors broadly disrupt proteostasis also suggests that they might activate HSF1 by an indirect mechanism, analogous to proteosome inhibitors (Gaglia, 2020)^30^.

On balance, we found evidence for functional Hsp90-HSF1 interactions convincing. Recently, Kolhe and co-workers (2023)^31^ mapped the Hsp90 interactome in yeast using site-specific photocrosslinking and clearly detected HSF1 interactions. Also intriguing, depletion of the co-chaperone Hsp70-Hsp90 organizing protein (HOP, or STIP1) was shown to significantly reduce HSF1 levels in human cell lines (Chakraborty, 2020)^32^. Lastly, two groups reported Hsp90 inhibitors disrupted Hsp90-HSF1 interactions *in vivo* using co-immunoprecipitation with Hsp90 variants (Kijima, 2018; Pesonen, 2021)^28,33^.

Our goal was to understand how molecular chaperones could achieve both feedback and forward regulation of HSF1. We saw key missing pieces to be: (1) molecular details of HSF1 activation and inactivation in a simplified system, (2) how Hsp90-HSF1 complexes form, and (3) the functional state of HSF1 in Hsp90 chaperone complexes. Here, using purified components and a multi-state FRET reporter system, we biochemically dissect HSF1 activation by heat and inactivation by Hsp40-Hsp70. To assess Hsp90-HSF1 interactions, we develop *in vitro* reconstitutions of Hsp90 variants containing an unnatural amino acid photocrosslinker, and with that system track both client and co-chaperone interactions. Side-by-side reconstitutions with HSF1 and the glucocorticoid receptor identify HSF1-specific requirements and chaperone mechanisms common between structurally divergent clients. Lastly, we use a combination of cryo-EM, structural, and biophysical methods to show that Hsp90 and HOP intercept HSF1 from Hsp70 and hold HSF1 stabilizing a monomeric intermediate activation state. Together, this work integrates decades of research into HSF1 regulation by molecular chaperones into a molecular description of the activation/inactivation cycle.

## Results

### Multivalent interactions of Hsp40-Hsp70 with HSF1 promote trimer disassembly

To begin, we developed a fluorescent assay to simultaneously monitor HSF1 conformational and oligomeric states. We wanted HSF1 constructs to report on the presence or absence of the auto-inhibitory coiled-coil interaction, as well as formation of the activating leucine zipper in the HSF1 trimer (or oligomer). In contrast to a previous result (Ahn, 2003)^34^, we found that cysteine-free HSF1(cf) was activated at the same temperature-concentration regime as wild-type HSF1 and bound to HSE-containing DNA olgionucleotides with the same affinity (Figure S1A). The C-terminal activation domain was not thought to affect HSF1 conformation, and we found truncated HSF1(1-417) behaved similarly to HSF1(wt) *in vitro*, as did a fluorescent protein fusion, HSF1(1-417)-GFP (Figure S1A). To monitor auto-inhibitory interactions, we generated HSF1(cf, L125C, 1-417)-GFP, with a cysteine incorporated in the region linking the DBD to Lz1-3, and labeled the single cysteine with maleimide-Alexa Fluor 546. In monomeric HSF1(cf,1-417, L125C-Alexa 546)GFP, the interaction of the autoinhibitory Lz4 with the Lz1-3 coiled-coil brings the GFP close to the Alexa546, allowing for Förster resonance energy transfer (FRET), whereas in the trimeric state FRET signal is greatly reduced (Figures 1B, S1B). To monitor oligomeric state, we paired the orange-green auto-inhibition reporter with a red-labeled HSF1 construct, HSF1(cf, L125C-alexa647), likewise conjugating maleimide-Alexa Fluor 647 into the DBD-Lz1-3 linker. In a mixture of free monomers, there is very little Alexa 546 to Alexa 647 FRET signal, but in the trimeric state the orange and red dyes are brought into close proximity (Figures 1B, S1B) (hereafter, HSF1(fret)). With our FRET reporter, we investigated how HSF1 dynamics are regulated by physiologically relevant biochemical components.

In cells, HSF1 activation is generally concomitant with nuclear import (Vujanac, 2005; Neueder, 2014)^35,36^. Depending on the reaction pathway for HSF1 oligomerization, local concentration could strongly affect HSF1 activation, and influence how HSF1 becomes activated by proteotoxic stressors other than heat. We examined the release of the Lz1-3-to-Lz4 auto-inhibitory interaction, and Lz1-3 oligomerization, by heating equal-molar mixtures HSF1(fret) at varied temperatures and concentrations. From initial rates, we determined HSF1 oligomerization at physiological temperature (37°C) and heat shock (41°C) is second order with respect to HSF1 concentration. Similarly, release of auto-inhibition is ∼2nd order at 37°C but lower order at higher temperatures (Figure S1C-E), demonstrating thermosensory function^23^. Thus in physiological and stress temperature ranges, we expect cellular HSF1 activation to be strongly dependent on local concentration, highlighting important modulatory roles for trafficking (nuclear import / export), degradation (e.g. by the proteosome), phase separation, and sequestration by molecular chaperones.

We evaluated how HSF1 is modulated by Hsp40 and Hsp70 chaperones and found that inactivation of trimeric HSF1 by Hsp40-Hsp70 is distinct from the unfolding of globular transcription factor clients. In agreement with a previous study (Kmiecik, 2020)^26^, we find that Hsp40 family member DnaJB1, Hsp70 (*HSPA1A or HSPA8*) and ATP-Mg are sufficient to displace HSF1(wt) trimers from DNA oligos containing a canonical HSE (Figure S2). Using the same HSP components and trimeric HSF1(fret), we saw addition of ATP led to a decrease in the oligomerization signal and a concurrent increase in auto-inhibition signal (Figure 1c). Notably, addition of DnaJB1-Hsp70-ATP to HSF1(fret) monomers does not affect the auto-inhibition signal, showing that the chaperones are not inactivating via unfolding HSF1 as seen with canonical substrates like p53, and GR. Curiously, during monomerization, we see the disappearance of the oligomer FRET signal faster than the appearance of the auto-inhibition signal. In contrast, during HSF1 activation by heat, release of auto-inhibition proceeds faster than oligomerization (Figure S3). These kinetics strongly suggest that the chaperone-mediated inactivation process proceeds through an open intermediate state, or states, that strongly favors HSF1 activation. In the presence of non-hydrolyzable ATP analogues, HSF1(tri)-DnaJB1-Hsp70 form very high molecular weight complexes (600-900kDa) (Figure S4). This demonstrated the necessity of ATP hydrolysis but also suggested binding of multiple copies of DnaJB1:Hsp70 to each HSF1 protomer.

Recently, studies of Hsp40-Hsp70-NEF systems with α-synuclein fibrils show that organization of multiple Hsp40:Hsp70s and coordinated ATP hydrolysis are critical to fibril disassembly by entropic pulling (Faust, 2020, Wentink, 2020; Beton, 2022; Monistrol, 2025)^37–40^, as also suggested for HSF1 inactivation. Synuclein disassembly requires an auto-inhibitory interaction in the J-domain of DnaJB1, absent in DnaJA1 and unnecessary for luciferase refolding (Faust, 2020)^39^. We likewise find that DnaJA1-Hsp70 is inactive with HSF1 trimers, and that the DnaJB1 H5 variant (J-domain always “on”) has low activity that decreases at high concentration. Likewise, removal of the DnaJB1 dimerization domain inhibits HSF1 monomerization (Figures S5A,B). In a simple competition assay, we added unlabeled HSF1(wt) trimers or monomers to a system of DnaJB1-Hsp70 inactivating HSF1(fret) trimers. HSF1(wt) trimers inhibited HSF1(fret) monomerization, whereas HSF1(wt) monomers had little effect (Figure S5C). Together, these results show that multiple Hsp40-Hsp70 binding events and coordinated ATP hydrolysis are critical for disassembly of HSF1 trimers.

### HspG0 and HOP slow the action of DnaJB1-Hsp70 on HSF1

To understand how Hsp90 regulates HSF1, we sought to incorporate Hsp90 and co-chaperones into our HSF1(FRET) system to evaluate their roles in the conformational kinetics of HSF1 or identify a stable intermediate state. Several co-chaperones have been suggested to regulate HSF1 alongside Hsp90, including p23, immunophillins (FKBP51, FKBP52), and HOP (Voellmy, 2007)^41^. Analogously, these interactions are well-characterized in the HSP-chaperone regulation cycle of the glucocorticoid receptor (GR) (reviewed, Deploey, 2023)^42^. We find addition of HOP-Hsp90 or HOP-Hsp90-p23 slows the rate of HSF1 inactivation relative to DnaJB1-Hsp70 alone; both the decrease in oligomerization signal (Fig. 1d) and increase in auto-inhibition signal (Figure S5d) are slowed. We note that addition of Hsp90 alone has a small effect. In contrast, addition of HOP alone, but not other co-chaperones, is sufficient to slow HSF1 inactivation (Fig 1d). Analogous trends were observed in the reverse process – heat shocking HSF1monomers co-incubated with chaperones – in which DnaJB1-Hsp70 slows trimerization at high temperatures but HOP and Hsp90 appear to lessen their effect. These data are consistent with HSF1 cycling though Hsp90 in complex chaperone mixtures, but we cannot rule out that the FRET signal was affected by sequestration of Hsp70 by HOP-Hsp90 or competition by HOP with DnaJB1 for the Hsp70 EEVD motif.

### Trapping HspG0-HSF1 using unnatural amino acid crosslinking

While previous studies had established that Hsp90 interacts with HSF1, we wanted to better understand what this interaction required and what it accomplished. To directly observe HSF1-Hsp90 interactions *in vitro*, we leveraged a site-specific crosslinking strategy and incorporated the unnatural amino acid *p*-benzoyl-L-phenylalanine (pBPA) (Chin, 2002)^43^ into recombinantly expressed human Hsp90. Upon UV (365 nm) exposure, the carbonyl carbon of pBPA rapidly and irreversibly inserts into nearby C-H bonds, thereby allowing one to trap protein-protein interactions at the pBPA interface. Sites for pBPA incorporation were chosen throughout *h*Hsp90AA, with a focus on the middle domain, and were guided by a recent *in vivo* crosslinking study (Kolhe, 2023)^31^ with yeast Hsp90 and our structural knowledge of several client interacting sites (Noddings, 2022; Wang, 2022)^44,45^ (Figure 2A).

**Figure 2:**
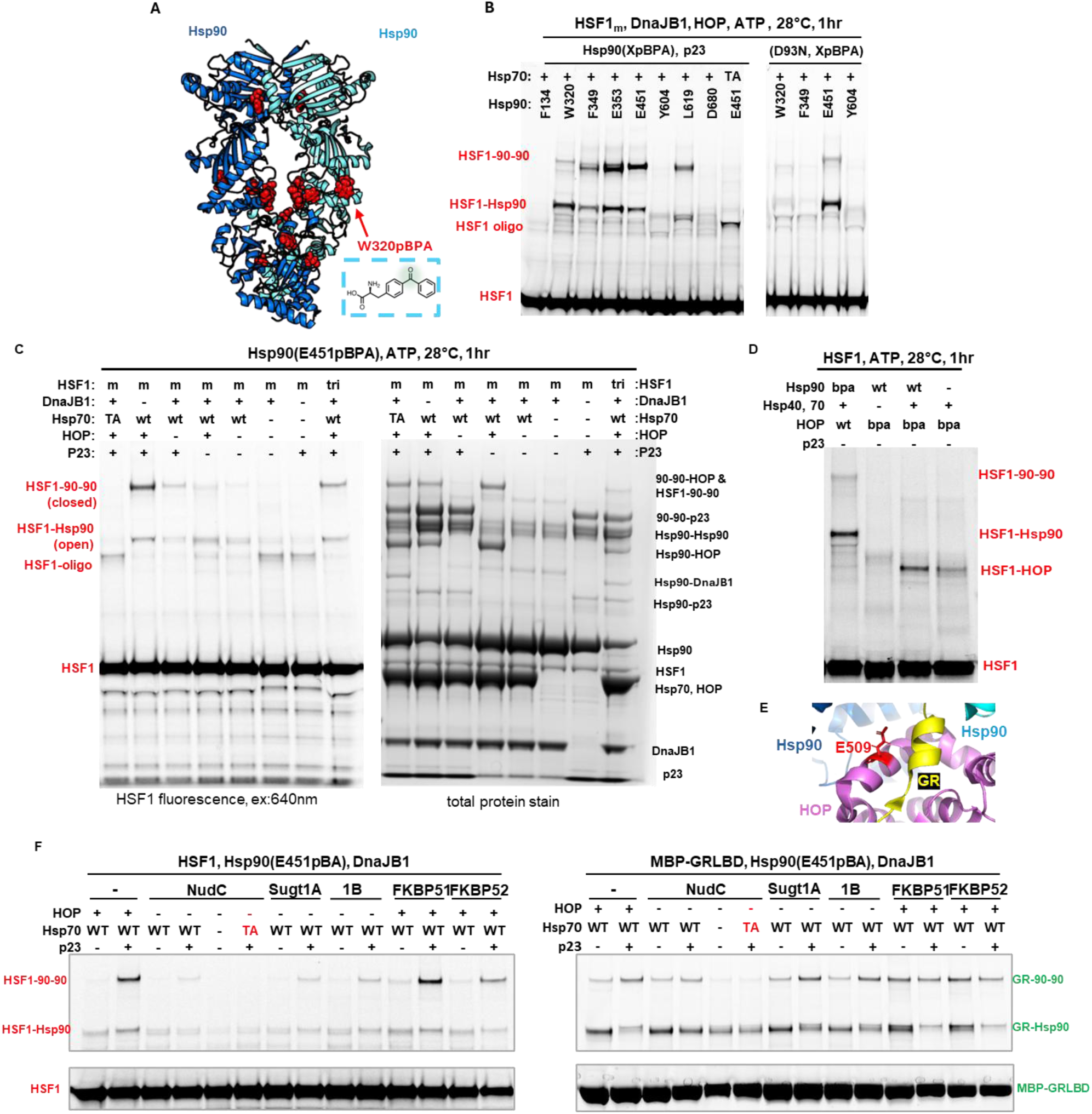
Functional Hsp70 and HOP promote HSF1-HspG0 complex formation. (A) Structure of human Hsp90AA (#7KRJ) dimer (blue, teal) shown in closed conformation. Amino acid positions substituted with pBPA are shown as red spheres. (B) DnaJB1 (3 µM), Hsp70 (12 µM), Hsp90 pBPA variants (12 µM), HOP (6 µM), and where noted, p23 (12 µM), were incubated with HSF1 monomers (4 µM) for 1 hr at 28 °C, then UV irradiated (365 nm) for 1 hr at 4 °C. ATPase deficient variants Hsp70(T204A) and HSp90(D93N) were used as noted. (C) HSF1(cf, Y225C-alexa647) monomers (m) or trimers (tri) were incubated with chaperones as in (B). Left, HSF1 and crosslinked adducts were imaged by in-gel fluorescence. Right, total protein was then stained by Coomassie. (D) HOP(E509pBPA) crosslinking to HSF1, conditions as (B) (E) GR (yellow) interacts with HOP (pink) at position E509 (red) in the structure of the GR-Hsp90 loading complex (#7KW7). (F) Side-by-side comparison of HSF1 (1.0 µM) and GR (1.0 µM) shows the effects of co-chaperones on Hsp90(E451pBPA)-client crosslinking. “GR” denotes MBP-tagged GR ligand binding domain, F602S, residues 521–777.

In initial experiments, we co-incubated pBPA containing Hsp90 variants, Hsp90(XpBPA), with HSF1, DnaJB1, Hsp70, HOP, p23 and ATP at 30 °C for 1hr, and then exposed these reactions to UV light at 4 °C. Using western blot detection, or in-gel fluorescence when Alexa 647-labeled HSF1 was included, we evaluated whether crosslinked Hsp90 species were generated. Encouragingly, we observed high-molecular weight HSF1 adducts in reactions that contained Hsp90 with pBPA positioned in or near the central cavity of the Hsp90 dimer (Figure 2A,B). Hsp90 is biased towards the closed (maturation) state in the presence of p23; we found that adding p23 with some Hsp90 pBPA constructs resulted in a second, higher adduct band. This band represents one of two HSF1-Hsp90 dimer species: both Hsp90 protomers crosslinked to the same HSF1 molecule, or one Hsp90 protomer crosslinked across the dimer interface, and the other Hsp90 protomer crosslinked to HSF1. We repeated some of these reactions using Hsp90 variants unable to bind ATP, Hsp90(D93N, XpBPA), that only adopt an open (loading) Hsp90 conformation. At some XpBPA positions, crosslinked species were much weaker (D93N-W320pBPA, D93N-F349pBPA), whereas Hsp90(D93N, E451pBPA)-HSF1 changes to a single crosslink (Figure 2B). This shift in crosslinking patterns thus provides information on state of Hsp90 when interacting with HSF1 in complex chaperone-co-chaperone mixtures. Notably, this result also demonstrates the utility of the photocrosslinking approach; previously, non-specific crosslinkers or mutations stabilizing the closed Hsp90 conformation were required to see significant Hsp90-HSF1 interactions by pulldowns (Guo, 2001; Kijima, 2018)^28,46^.

Many paradigms of Hsp90-client interactions are known, and we sought to identify which were possible with HSF1-Hsp90. Whereas some Hsp90 clients require specific co-chaperones to bring them to Hsp90, others are promiscuous, and others may associate with Hsp90 directly (Silbermann, 2024)^47^. Recent studies have highlighted that Hsp90 clients tend to contain intrinsically disordered regions (IDRs) (Babu, 2024)^48^, and the well-known client human Tau is both IDR-rich and associates with Hsp90 directly (Karagoz, 2014)^49^. Although HSF1 contains large IDRs, we find Hsp90-HSF1 interaction requires Hsp40 and Hsp70 for minimal signal and HOP for strong association (Figure 2C). Intriguingly, in a cycling chaperone system at low temperature (28 °C), Hsp90-HSF1 crosslinks are seen whether the reaction is begun with monomeric or trimeric HSF1. Under these conditions, we do not see significant monomer to trimer conversion in our HSF1(fret) assays, likely because DnaJB1-Hsp70 activity is fast relative to trimerization. Nonetheless, inactivation of Hsp70 by T204A mutation (ATPase dead) completely prevents HSF1-Hsp90 crosslinking.

Total protein staining from these experiments provided unexpected insight into the interplay among Hsp40-Hsp70-Hsp90-HOP-p23. With components expected to generate an Hsp90 maturation state, strong bands are seen for Hsp90-Hsp90-p23, showing that the disordered p23 C-terminal tail likely reaches into the Hsp90 dimer lumen (Figure 2A). In a system with just Hsp90-HOP-p23 (Figure S6A), HOP prevents p23 induced Hsp90 closure, and active Hsp40-Hsp70 are required to fully reach the Hsp90 maturation state. In available structures of closed Hsp90, E451pBPA is positioned to crosslink across the dimer interface, but would not be expected to form Hsp90-Hsp90 crosslinks in open Hsp90 conformations (Noddings, 2022, Wang, 2022)^44,45^. Nonetheless, Hsp90-Hsp90-HOP crosslinks are seen in a two-component system, corroborating that HOP stabilizes a partially closed state (Southworth, 2011)^50^. Lastly, we were surprised to see low-levels of Hsp90-DnaJB1 crosslinking, however, eukaryotic cells can overcome HOP deletion by forming bacterial-like Hsp40-Hsp70-Hsp90 complexes (Bhattacharya, 2020)^51^, consistent with our observed low levels of HSF1-Hsp90 interaction in the absence of HOP.

We wondered whether HOP interacts directly with HSF1 to promote its loading into Hsp90, as recently demonstrated for HOP and Hsp90 client GR in the Hsp90 loading complex (Wang, 2022)^45^. We purified HOP(E509pBPA) and incorporated it into our reconstituted system. HOP(E509pBPA) with only Hsp90(wt) does not crosslink HSF1, DnaJB1-Hsp70-HOP(E509pBPA) allows for weak crosslinking, and a full loading state system with Hsp90 results in stronger crosslinking (Figure 2d). Deletion of the DP2 domain weakens Hsp90-HSF1 crosslinking at multiple positions in both loading and maturation state systems, suggesting that the weakened interaction is not due to conformational shifts (Figure S7B,C) but to inefficient complex formation. We envision that HOP helps to pre-organize Hsp70-bound HSF1 for loading onto Hsp90, as has been proposed for GR, but with a stronger HOP-dependence for HSF1 in the same reconstituted systems (Figure S7C).

Although core chaperone machinery is strongly conserved across eukaryotes, co-chaperone levels vary by cell type while providing semi-redundant pathways into the Hsp70 and Hsp90 cycles (Genest, 2019; Johnson, 2021)^52,53^. HOP, for example, can be knocked out in human cell lines (Bhattacharya, 2020)^51^ but not in a mouse, and co-chaperones like Sugt1 (Engler, 2026)^54^ and NudC (Biebl, 2022)^55^ provide alternate Hsp90 client loading pathways. Of particular note, NudC was recently shown to transfer both GR and p53 to Hsp90 from Hsp40-Hsp70 utilizing a new pathway. We performed side-by-side reconstitutions of HSF1-Hsp90 and GR-Hsp90 interactions in a panel of cochaperone mixtures (Figures 2D, S7A). In contrast to GR, neither NudC nor Sugt1 can substitute for HOP in promoting HSF1-Hsp90 interaction. FKBP51 and FKBP52 modestly alter HSF1-Hsp90(E451pBPA) crosslinking strength, however, they may do so by promoting a closed Hsp90 state over a loading state and also cannot substitute for HOP (Figure S7B).

The inability of Sugt1 and NudC to substitute for HOP was surprising given that they are both dimers and therefore could provide multiple client interacting sites in a manner analogous to HOP-Hsp702. Moreover, as with HOP, we observe Hsp90(E451pBPA)-NudC, Hsp90(E451pBPA)-Sugt1, and 90-90-NudC/Sugt1 crosslinks (Figure S7A). Nonetheless, it is clear from our data that these two co-chaperones operate via distinct mechanisms. NudC appears to inhibit closure of Hsp90 and competes with p23 (reduced Hsp90-Hsp90 and 90-90-p23 crosslinking, Figure S7A), and p23 does not promote GR ligand binding in an Hsp40-Hsp70-NudC-Hsp90 system (Biebl, 2022)^55^. The cycle of Hsp90 with NudC may therefore be distinct from the classical HOP-mediated loading-maturation cycle in a way that alters or prevents HSF1 association with Hsp90. In contrast, we see strong Sugt1-mediated GR-Hsp90 interactions that are p23-modified (Figure S7a) but only weak HSF1-Hsp90 interactions. We propose that the differences between GR and HSF1 in these systems are that HSF1 requires direct transfer from Hsp70, and that HSF1-HOP interactions are additionally stabilizing of intermediate HSF1 states.

### High temperature and inhibitors push HspG0-HSF1 into the loading state

To clarify the role of Hsp90-HSF1 interactions, we tested how they responded to stresses that activate HSF1. First, we reconstituted DnaJB1-Hsp70-HOP-Hsp90-p23-HSF1 systems as time course experiments across a range of temperatures (Figure S8). At higher temperatures, both Hsp90-HSF1 (typically open state) and 90-90-HSF1(closed state) crosslinks reached peak intensity faster than at 30 °C, reflecting faster cycling of the system^56^. Crosslink intensity, however, was much stronger at 37 °C than 42 °C for both states at multiple pBPA positions (Figure S8A-E). Although this could be due to HSF1 oligomerization “winning” over Hsp90 association, a similar difference between 37 °C and 42 °C was seen with Hsp90-GR crosslinking (Figure S8f). In total protein staining we observed correspondingly weaker Hsp90-p23 and Hsp90-Hsp90 crosslinking, suggesting that the system was shifting towards an open Hsp90 state (Figure S8d,e). We used HOP(E509pBPA) + Hsp90(W320pBPA) system to look at loading states specifically, and observed both higher temperatures and longer time points promoted HSF1-HOP-HSF1 accumulation (Figure S8G,H). High temperatures biasing the system bias towards open Hsp90 might allow faster disassociation of HSF1 in response to elevated heat shock temperatures, either by exchange in and out of the loading state, or by faster cycling through the closed state. The speed of HSR induction varies, but translocation of HSF1 from the cytosol to the nucleus occurs over minutes (Baler, 1993; Neueder 2014)^36,57^, as do the most rapid transcriptional changes in response to heat shock (Mahat, 2016)^3^. Notably this is a similar time scale to cycling of HSF1 through the closed Hsp90 states at 42 °C (Figure S8A,C,D).

Activation of the heat shock response by Hsp90 inhibitors remains a serious impediment to therapeutic development, and we sought to clarify how they affected Hsp90-HSF1 interactions. In a maturation state system, increasing concentration of the N-terminal Hsp90 inhibitor AUY-922 blocks 90-90-HSF1 and Hsp90-p23 crosslinking, expected with inhibition of ATP binding and Hsp90 dimer closure. Likewise, AUY-922 blocked 90-90-GR crosslinking, while both Hsp90-HSF1 and Hsp90-GR bands were strengthened, suggesting an Hsp90 loading state (Figure S9a,b). It thus follows that in a forced loading state system Hsp90(D93N, W320pBPA)-HOP(E509pBPA)-HSF1 crosslinking was constant with increasing AUY-922 concentration (Figure S9c). Similar effects were observed with an alternate N-terminal Hsp90 inhibitor, XL-888, and with increasing temperature (Figure S9d). For many Hsp90 clients, Hsp90 inhibition leads to their ubiquitination (on or off Hsp90) and degradation via the proteosome. In contrast, activation of HSF1 by Hsp90 inhibitors occurs without notable degradation, and is slow compared to heat shock (Kijima, 2018; Gaglia, 2020)^28,30^. Thus, comparison of heat shock and Hsp90 inhibition suggests they may induce the HSR via different pathways.

Hsp90AB-specific inhibitors promote degradation of some clients without inducing the HSR (Dernovšek, 2023; Reynolds, 2025)^58,59^, suggesting that heat-inducible Hsp90AA and constitutive Hsp90AB may differentially regulate HSF1. We compared HSF1 and GR interactions with Hsp90AA(E451pBPA) and Hsp90AB(E443pBPA) under loading and maturation state conditions (Figure S10). For both clients, the interactions with the two Hsp90 isoforms appeared similar, and as with Hsp90AA, ATP-competitive inhibitors block Hsp90AB-p23 crosslinking and maturation state formation. A major difference, however, is seen with increasing AUY-922 concentration at 30 °C vs 42 °C. At 30°C the inhibitor reduces Hsp90AB-HSF1 crosslinking but at 42 °C the inhibitor strengthens in interaction in a maturation system (Figure S11A,B), whereas it decreases at 42 C with Hsp90AA. In the same conditions the loading state Hsp90AB(E443pBPA)-HOP(E509pBPA)-HSF1 crosslink is strong and constant (Figure S11C,D). These results suggest the ATP-competitive inhibition of Hsp90AB induces a state with higher affinity for HSF1 than the same treatment of Hsp90AA.

### Structural characterization of HspG0-HSF1 complexes

Using cryo-EM, we next sought to resolve the interactions between Hsp90 and HSF1 and solved several structural states of Hsp90-HSF1. To do this, we reconstituted HSF1-chaperone complexes that were stabilized by pBPA photocrosslinking. This avoided the need for stabilization by glutaraldehyde as required for GR and kinase structures, greatly facilitating analysis. Free chaperone components were removed afterwards in additional purification steps. To our knowledge, this strategy has been used only once before (Xie, 2025)^60^. As a general principle for multi-protein samples, we recommend minimizing both the number of pBPA sites, and choosing sites with high crosslinking efficiency and only one possible interacting partner.

To resolve the Hsp90-HSF1 maturation state, we set up reaction conditions that contained monomeric Twin-Strep-HSF1(wt), Hsp90(E451pBPA), DnaJB1, Hsp70, HOP and p23, and ATP-NaMO4. Crosslinked HSF1-Hsp90-co-chaperone complexes were isolated via Strep-Tactin resin, concentrated, and frozen on grids (Figure S12). We obtained a consensus Hsp902-HSF1-p23 structure at ∼2.7 Å with clear density for HSF1 threaded through the Hsp90 dimer lumen (Figure 3A). The E451pBPA crosslink is primarily made to R355 on the opposite Hsp90 promoter, however, there is a mixture of states and crosslinking to HSF1 is also present. Focused 3D classification on p23 and the Hsp90 NTDs showed states with zero or one copy of p23 bound but with the Hsp90s similarly closed. We used the asymmetric class with p23 for additional 3D classification on the Hsp90 MDs and bound HSF1 and resolved at ∼2.9 Å a state with clearer crosslinks and HSF1 (Figures 3B, S12). Additional work (below) was used to guide assignments of HSF1 residues bound to Hsp90.

**Figure 3:**
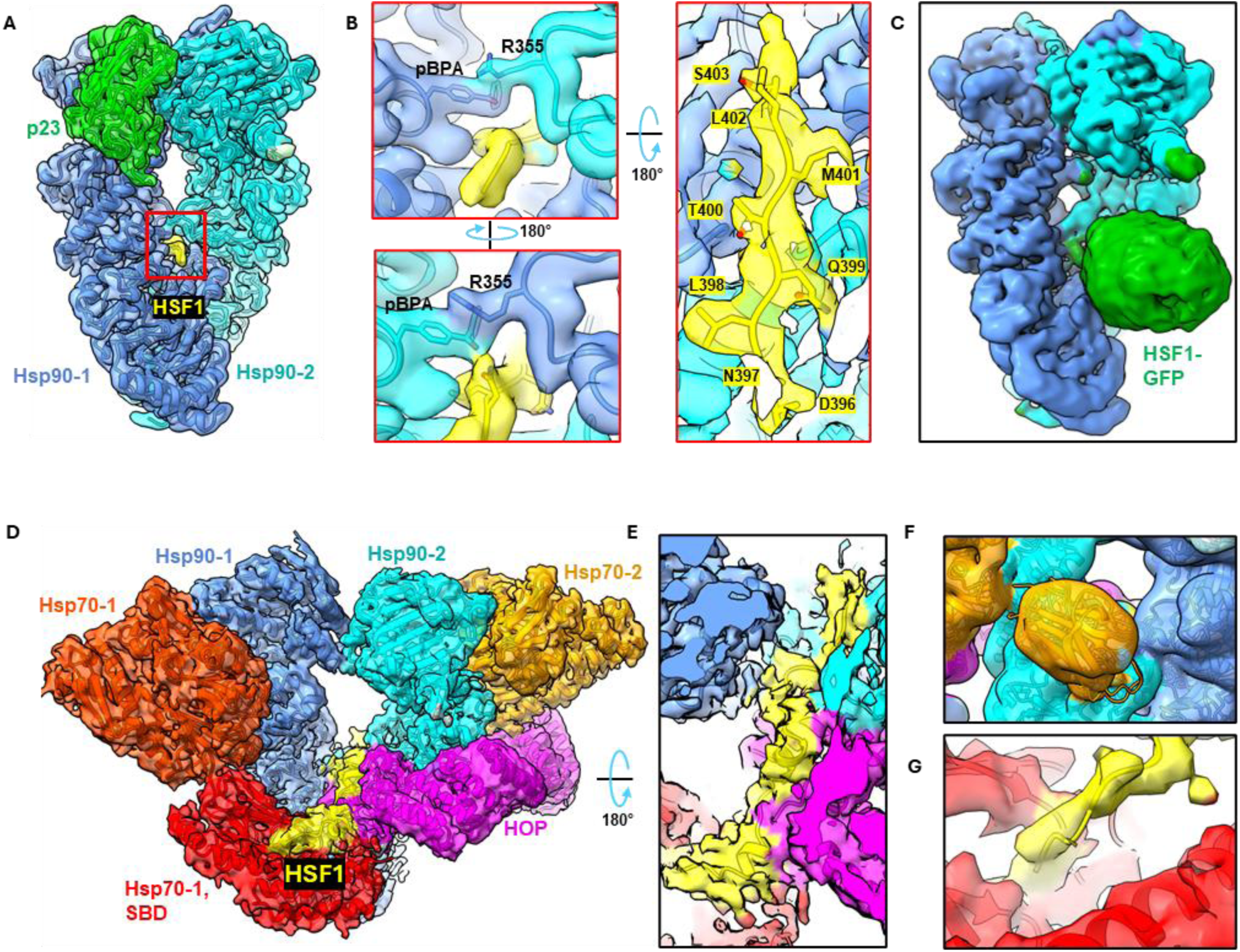
Cryo-EM structures of HSF1-HspG0 complexes show extensive, conserved contacts. (A) Maturation state Hsp90 complex with HSF1 (yellow), Hsp90(E451pBPA) (blues), and p23 (Green). The red box highlights views in (b). (B) At left, pBPA 451 and R355 are shown as sticks in view of both sides of the Hsp90 lumen. At right, client HSF1 is shown from above with sidechain assignments. (C) HSF1(1-417)-GFP (green) is trapped in a closed Hsp90 dimer. (D) Loading state HSp90 complex with, HSF1 (yellow), Hsp90 (blues), Hsp70 (oranges), HOP(E509pBPA) (pink). (E) Top, cut-away view shows HSF1 contacts with Hsp90, HOP, and Hsp70-1. (F) Low-pass filtered map showing the location of the β-domain of the Hsp70-2 SBD, on the face of the Hsp90 dimer opposite HOP. (G) HSF1 passes through the lumen of the Hsp70-1 SBD adjacent to HOP.

In parallel, we obtained a closed Hsp902-HSF1(1-417)-GFP structure using the same workflow as with HSF1(wt). To our surprise, modest density for the GFP was present adjacent to the Hsp90 MDs and luminal HSF1 (Figure 3C, S12). The region linking the luminal HSF1 to the GFP is too low resolution to model, but the arrangement nonetheless strongly suggests that the HSF1 Lz4 domain or flanking residues are bound by Hsp90. Although p23 was also present in the initial consensus map, focused classification on the GFP and Hsp90 MDs separated states with either p23 or GFP present. Possibly, the GFP is displacing p23 and ADP-MO4 keeps Hsp90 closed. Notably, reconstitution of the HSF1-Hsp90 interactions with tailless p23(1-112) still generates a maturation state with strong Hsp90(E451pBPA)-HSF1 crosslinking (Figure S12). Thus, in contrast to the Hsp90-GR system in which there is a specific p23 tail helix interaction with GR, we conclude the primary role of p23 in the HSF1 chaperoning is to promote Hsp90 closure.

To visualize loading state complexes (HSF1-Hsp902-Hsp702-HOP), we set up reactions with monomeric HSF1(wt), Hsp90(D93N), DnaJB1, Hsp70, and HOP(E509pBPA). Crosslinking reactions were separated by size exclusion chromatography, peak fractions were analyzed by SDS-PAGE, and those containing crosslinked HSF1 were concentrated and frozen on grids (Figure S13). Although HSF1 has a low molecular weight relative to the chaperone components of this complex, HSF1 has a very large hydrodynamic radius, especially as a trimer. The clear separation of HSF1 containing complexes from empty loading states (Figure S13) suggests that the client HSF1 may be in an extended conformation (see below). Ultimately, we obtained an ∼3.2 Å reconstruction in which all six proteins are present; individual components were better resolved by focused classification and local refinement (Figures 3D, S13).

By comparing the structures of the HSF1 and GR loading complexes, we observed several new features despite their similar general architectures. The Hsp90 dimer adopts a semi-closed conformation in both with a large space between the middle domains, and the NTDs are touching but not crossed over. Two copies of Hsp70 are present, with their nucleotide binding domains (NBD) making identical and extensive contacts to the Hsp90 NTDs and MDs. ADP is bound in each Hsp70 NBD, whereas Hsp90 is nucleotide-free (Figure S13G). Previous work showing these Hsp70-Hsp90 contacts accelerate Hsp90 ATPase (Genest, 2015)^61^ is also reflected in the better ability of Hsp70(wt) to promote the maturation state than Hsp70(T204A) (Figure 2a). HOP(E509pBPA) is situated as in the GR-Hsp90 loading complex, with TPR1B, TPR2, and the DP2 domain resolved, collectively wrapping around one Hsp90 protomer but making contacts with all the components.

We observe HSF1 density threaded through the lumen of Hsp90 and contacting the HOP DP2 domain. Intriguingly, a short helical segment is present at the HSF1-HOP interface (Figure 3e), as seen with GR, and may be indicative of a client preference of HOP. The predicted HSF1 structure is largely disordered outside of the DBD and Lz1-3 domains, however, several short helical segments are predicted near the Lz4 helix. With HOP E509pBPA, low map quality likely reflects multiple crosslinking modes suggesting heterogeneous binding to HOP. Despite extensive classification and refinement, we were unable to unambiguously model specific HSF1 residues into the loading complex density. The tube of HSF1 density is continuous with Hsp70 SBDs on each side of the Hsp90 MDs, one contacting HOP DP2 and the opposite somewhat free (Figure 3E,F). Notably, this architecture extends the region of client HSF1 occluded in the loading complex. In contrast, only one Hsp70 SBD was seen in the GR loading complex (Wang, 2022)^45^, but both our crosslinking results and reconstitutions of HSF1-DnaJB1-Hsp70 systems (Figures S4, S5) suggest that several Hsp70 may bind HSF1 simultaneously.

The Hsp70 SBD adjacent to HOP DP2 is also mobile but can be well-resolved by focused 3D classification and refinement. The primary mode of movement is rotation around an axis parallel to HOP (Figure S13F). At one end of the rotation, only the Hsp70 SBD-β domain is resolvable, analogous to the GR loading structure. At the other end of the rotation, the full SBD is resolved, with the SBD lid clamped closed on the SBD-β. Continuous density for an unstructured region of HSF1 passes from the HOP DP2 through the center of the SBD binding pocket (Figure 3G). Altogether, an extended region of HSF1 is bound in this state. The disordered regions of HSF1are relatively hydrophobic, characteristic of both HOP and Hsp70 substrates, and contain Hsp70 binding sites at 347-375 and 442-471 (Kmiecik, 2020)^26^. Additionally, PTMs at several sites in HSF1 have been proposed to modulate its interactions with Hsp90 (Gomez-Pastor, 2018)^2^.

### HspG0 binds at or adjacent to LZ4

To better identify the Hsp90 and HOP interaction sites in HSF1 we used photocrosslinking in combination with mass spectroscopy. In loading state conditions, we detected HOP(E509pBPA)-HSF1 crosslinks in four clusters with adjacent or overlapping Hsp90(E451pBPA)-HSF1 crosslinks at three clusters (Figure 4A). These include sites in the trimerization domain and at the C-terminus of the activation domain, but the preponderance of detected adducts were found within residues 380-430, containing the auto-inhibitory helix and adjacent regions. In contrast, in the maturation state, Hsp90(E451pBPA)-HSF1 crosslinks were detected only within HSF1 residues 380-407. This matches residues 396-403 directly observed in the maturation state cryoEM structure (Figure 3B). Although these results do not necessarily identify all interaction sites, and may be somewhat biased by differential fragment ionization, the protein digest conditions chosen allowed detection of nearly all of HSF1 when analyzed alone. These results imply some conservation of the Hsp90-client mechanism observed with GR, despite differences in client structure, in which loading and maturation state client interaction overlap but unstructured client regions may slide through the Hsp90 lumen.

**Figure 4:**
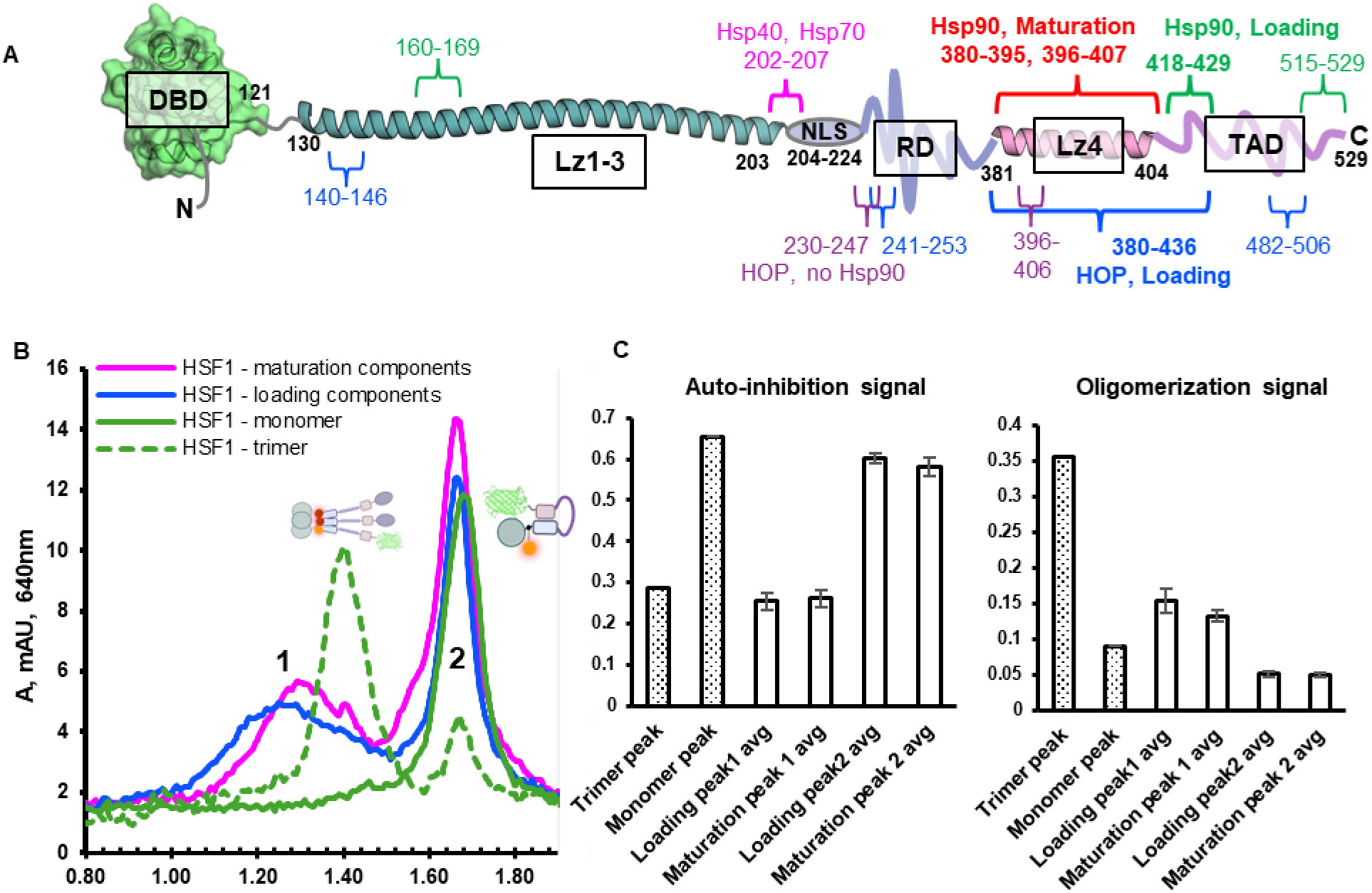
HspG0 binding to HSF1 stabilizes an extended monomeric conformation. (A) This cartoon summarizes the results of crosslinking-MS experiments with HSF1. Hsp90(E451pBPA)-HSF1 adducts in loading complexes (green) and maturation complexes (red), and HOP(E509pBPA)-HSF1 adducts in loading complexes (blue) were separated from free components by analytical SEC. Fractions containing adducts were then concentrated, digested, and analyzed by LC MS-MS. (B) Analytical size exclusion chromatography was used to compare and separate HSF1(fret) alone to from HSF1(fret) in chaperone complexes. (C) FRET signal was analyzed from peak fractions 1 and 2 shown in (B). Hsp90-HSF1(fret) complexes prepared this way are not auto-inhibited and appear largely monomeric; free trimeric HSF1(fret) may have been incompletely separated.

### HspG0 stabilizes an extended form of monomeric HSF1

To further probe the state of HSF1 in complex with Hsp90, we isolated Hsp90-HSF1 complexes reconstituted using the HSF1(fret) system. As previously, loading and maturation state Hsp90(W320pBPA)-HSF1 were separated over SEC and different HSF1 species located by fluorophore absorbance (Figure 4B). Monomeric and trimeric HSF1(fret) were run separately for comparison. In both Hsp90 systems, two HSF1 peaks were observed over SEC, one with the same retention as monomer and the other running ahead of trimeric HSF1. Pooled fractions from peak 2 had the same auto-inhibition and oligomerization FRET signal as monomers; in contrast pooled fractions from peak 1 (Hsp90 complex fractions) were predominantly monomeric but not auto-inhibited. Together, these results suggest that Hsp90 maintains HSF1 in an extended state rather than participating directly in disassembly of HSF1 trimers. Contrary to most previous reports, Hsp90 binding to HSF1 Lz4 could potentiate HSF1 activation by blocking autoinhibition.

We sought to directly evaluate if Hsp90 affects HSF1 trimerization. We prepared photocrosslinked Hsp90-HSF1 complexes, then added twinStrep-HSF1 (bait) to those reaction mixtures, raised the temperature to allow trimerization with Hsp90-HSF1 complexes (prey), and quenched the reactions on ice for a subsequent pulldown using Strep-Tactin resin (Figure 5A,B). In parallel loading, maturation, and (-)Hsp90 conditions, little FLAG-HSF1 (prey) was pulled down as free HSF1 or as adducts following room temperature incubation. Increasing incubation temperature did not change the amount of HSF1 bait pulled down but dramatically increased the amount of prey HSF1 in the Strep-Tactin eluates (Figure 5B). Both uncrosslinked HSF1 and Hsp90/HOP-HSF1 adducts were pulled down, demonstrating that they had formed mixed trimers with twinStrep-HSF1. Trimerization of Hsp90-HSF1 adducts with free HSF1 was observed independent of pBPA site (W320, E353, or E451) and with Hsp90AB as well as Hsp90AA (Figure S14). Densitometry analysis comparing the adduct/free ratio of prey HSF1 suggests that at lower temperatures HSF1-Hsp90 adducts are enriched relative to free HSF1 (Figure S14C). Taken together, these results point to Hsp90 binding to and occluding the auto-inhibitory domain of HSF1, thereby potentiating re-trimerization.

**Figure 5:**
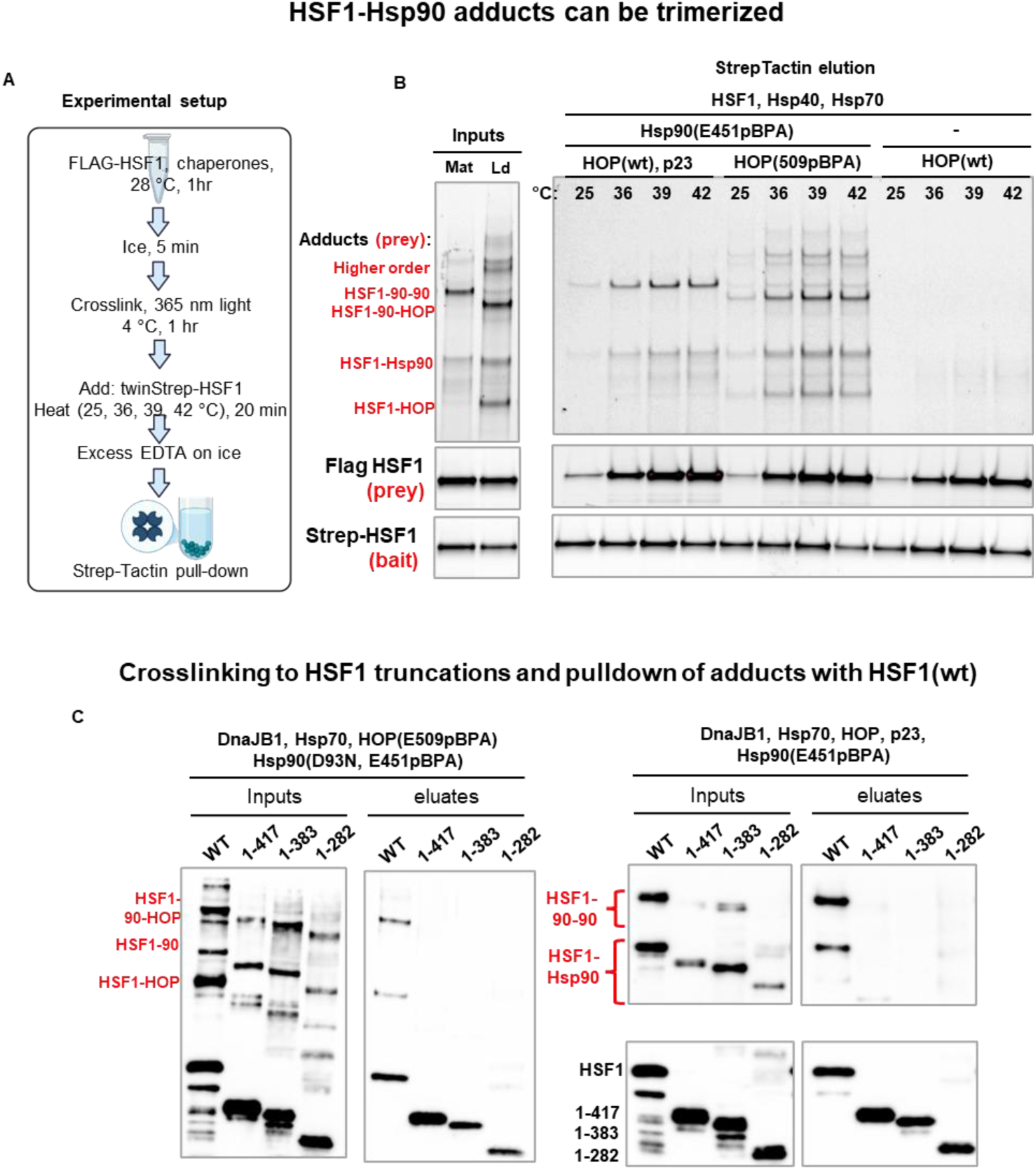
HSF1 bound to HspG0 can trimerize, but secondary binding sites inhibit trimerization. (A) Experimental setup: prey crosslinked HSF1(cf, Y225C-alexa647)-Hsp90 complexes were heated with bait twinStrep-HSF1(cf, Y225C-alexa546) and trimerization of HSF1-Hsp90 adducts assessed by streptavidin pulldown. (B) Inputs (left) show prey HSF1-chaperone adducts prior to sample heating, and pulldowns (right) show what was pulled down by bait HSF1 following heating at the indicated temperatures. The right panel compares maturation (lanes 1-4), loading (lanes 5-8), and Hsp90-free (lanes 9-12) HSF1-chaperone complexes. (C) Bait-prey experiments were performed as in (B) using FLAG-HSF1(wt) and a series of HSF1 truncations. These truncations force Hsp90 binding to HSF1 at secondary sites N-terminal to HSF1 Lz4, and this blocks the trimerization of HSF1-Hsp90 adducts with free, wild type HSF1.

Although Hsp90-HSF1 interaction at HSF1 Lz4 appears favored, our crosslinking-MS analysis (Figure 4A) identified additional interaction sites. To validate and understand these interactions, we tested Hsp90(E451pBPA) and HOP(E509pBPA) crosslinking to a series of HSF1 truncations at the activation domain (HSF1(1-417)), auto-inhibitory domain (HSF1(1-383)), or within the regulatory domain (HSF1(1-283)). As described above, we performed bait-prey pulldowns and then detected free and crosslinked FLAG-HSF1 by western blot (Figure 5C). Firstly, the total amounts of crosslinked products appear lower for the HSF1 truncations, particularly HSF1(1-282) (Figure 5C, “inputs”). Also notable is that crosslinking of the truncations to HOP(E509pBPA) is reduced in loading state conditions, as is the formation Hsp90-Hsp90-HSF1 adducts in maturation state conditions. This again suggests that the dominant mode of Hsp90-HSF1 interaction is at Lz4. Roughly equal amounts of free WT and truncated HSF1 are present in the eluates under all conditions (Figure 5C, “eluate”). In contrast, very little or none of the truncated crosslinks are pulled down, consistent with HSF1-Hsp90 interactions forced to be biased towards interacting in the trimerization domain (Figure 4A). In summary we observe two types of Hsp90-HSF1 interactions; a primary mode of binding at HSF1 Lz4 that promotes trimerization of HSF1, and secondary modes in other domains of HSF1, some of which inhibit trimerization.

### Trimerized HspG0-HSF1 complexes can bind heat shock elements

Hsp90 might still negatively regulate HSF1 by preventing it from binding its target promoters, as one report previously suggested (Kijima, 2018)^28^. We prepared photocrosslinked Hsp90-HSF1 complexes, heat-shocked them in active chaperone mixtures, then quenched them on ice and briefly incubated them with HSE-containing DNA oligonucleotides. To evaluate whether DNA was bound to HSF1, we separated these mixtures by SEC and tracked both the DNA and HSF1 by fluorescent tags (Figure 6A,B). Canonical HSEs remained bound to heat-shocked (trimerized) HSF1-Hsp90 maturation complexes, as seen with HSF1 trimers alone, as did DNA oligos containing the main HSE in the *HSPSOAA* promoter (HSE90AA). Addition of excess Bag1 did not affect the retention of HSE-Hsp90-HSF1, showing that Hsp70 in the mixture was not responsible for the shift relative to HSF1 trimers. The *HSPSOAA* promoter HSE is structured to allow binding of two HSF1 trimers, and concordantly, there is a large shift in retention volume of the HSE90AA-Hsp90-HSF1 sample (Figure 6A,B). To show that DNA binding by Hsp90-HSF1 complexes rather than free HSF1 was responsible for the retention shifts, we analyzed fractions from these SEC experiments by SDS-PAGE and in-gel fluorescence (Figure 6C,D). In two states, Hsp90(E451pBPA)-HOP(509pBPA)-HSF1 (loading) and Hsp90(W320pBPA)-HSF1 (maturation) adducts were shifted to a shorter retention volume by heat shock and then further shifted by addition of HSE90AA. From these results we conclude that HSF1 trimers can bind DNA even when integrated into Hsp90 complexes.

**Figure 6:**
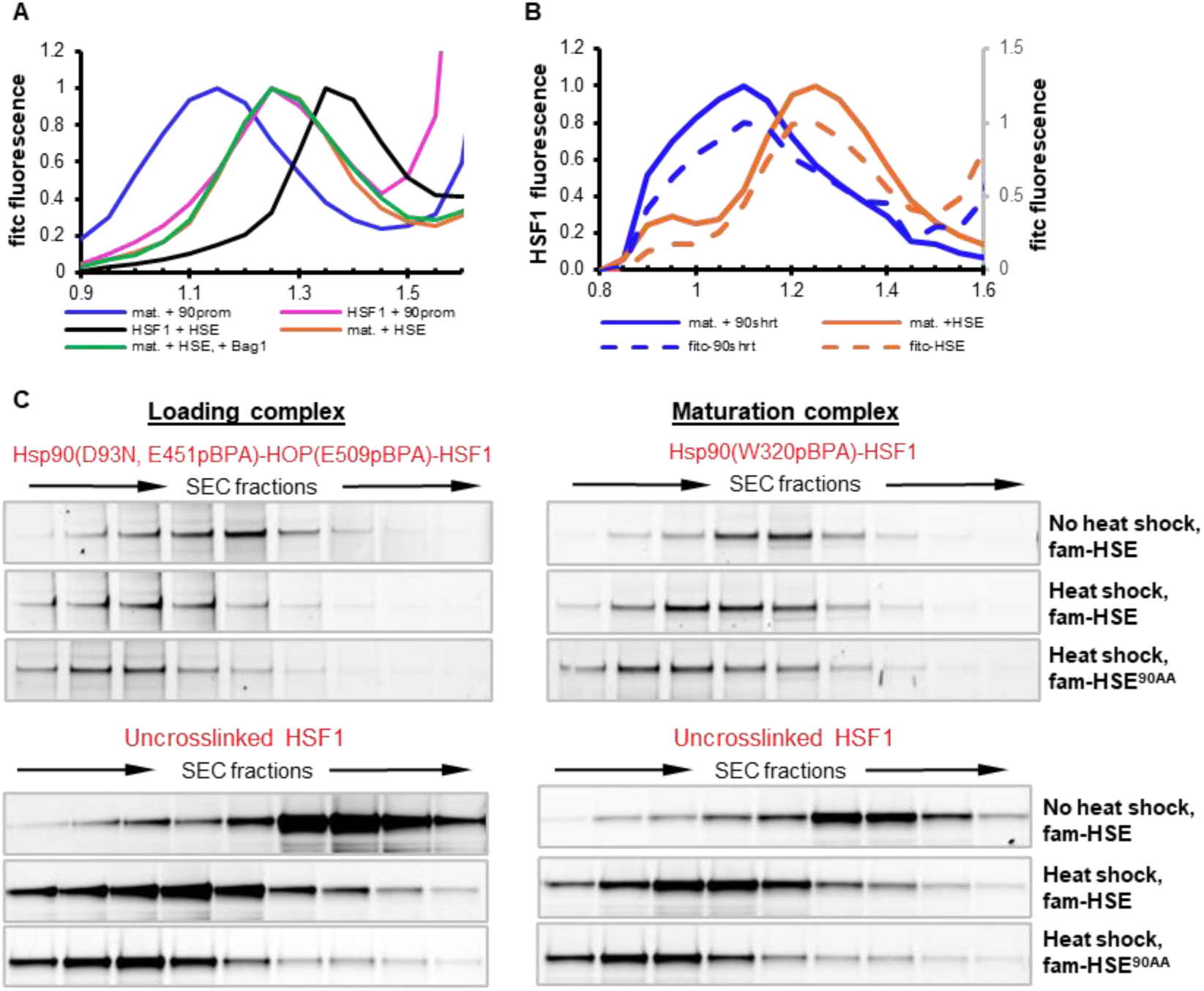
HSF1 trimers containing HSF1-HspG0 complexes bind stably to heat shock elements. (A) HSF1 and chaperones to promote maturation state complexes were prepared, photo-crosslinked for stabilization, heat shocked to promote trimer formation, then quenched and incubated with fitc-labeled DNA oligos on ice, either canonical HSEs or the Hsp90AA promoter HSE90AA. These mixtures were then separated by analytical size exclusion chromatography. (B) Experiments were prepared as in (A); a mixture of HSF1(wt) and HSF1(cf, Y225C-Alexa546) was used to allow simultaneous tracking of HSF1 and the labeled oligos. (C) HSF1-chaperone-DNA complexes were separated as in (B), using loading state (left) or maturation state (right components). Comparison of free HSF1 and crosslinked HSF1-chaperone adducts shows they have similar retention shifts with heat shock and addition of DNA.

## Discussion

Here, we pursued a detailed biochemical and structural study of HSF1 interactions with the Hsp40-Hsp70-Hsp90 chaperone system to provide mechanistic insights on HSF1 dynamics in cells. We initially hypothesized that chaperones regulate transcription factor clients in a universal manner shared by HSF1, p53 and steroid hormone receptors. HSF1 is structurally very distinct, however, and we found important mechanistic differences in chaperone-HSF1 regulation. In particular, we found that Hsp40-Hsp70 inactivates trimer HSF1 by physically separating its monomers instead of unfolding HSF1 into an aggregation-prone state (GR: Kirschke 2014; Moessmer 2022, p53: Dahiya 2019, Boysen 2019)^62-65^ (Figure 1B-D). Biochemically, inactivation of trimeric HSF1 by Hsp40-Hsp70 more closely resembles disaggregation of fibrils or amyloids with multiple Hsp70-40s being involved. By this analogy, we thought Hsp90 might not maintain HSF1 in a near-native state as is the paradigm for Hsp90 folding clients66.

Unlike previously well-characterized clients, several conflicting models have been proposed for Hsp90’s role in HSF1 regulation. Variously, these were that Hsp90 has little effect, that Hsp90 holds HSF1 monomers in a repressive complex, potentiates HSF1 activation, or removes trimeric HSF1 from DNA. Our results allow us to define a rationale for these conflicting models by unambiguously demonstrating key Hsp90-HSF1 interactions that are consistent with multiple regulatory regimes depending on the cellular context.

Together, our multi-pronged biochemical, structural, and MS analyses show that the full Hsp40-Hsp70-Hsp90 chaperone system interacts with HSF1 in several modes (Figure 7). In a simple system, heat stress trimerizes and activates HSF1, then Hsp40-Hsp70 monomerizes and inactivates HSF1 as stress subsides (Figure 7A). The presence of Hsp90 and co-chaperones allows for regulation throughout the HSF1 activity cycle (Figure 7B). Hsp40-Hsp70-HOP can bring HSF1 into Hsp90 loading complexes without significant accumulation of trimeric HSF1 (Figure 7B ‘cytoplasm’, Figure 2C). Alternatively, as Hsp40-Hsp70 disassemble HSF1 trimers, HOP may intercept the process and bring HSF1 to Hsp90 (Figure 7B ‘nucleus’). Predominantly, Hsp90 binds HSF1 near the Lz4 auto-inhibitory domain in a manner that stabilizes an extended, pre-activated state that potentiates HSF1 trimerization (Figures 4, 5). The same extended HSF1 monomeric intermediate on Hsp90 can efficiently relax back to an auto-inhibited monomer. Thus, Hsp90 interactions enhance the lifetime of this poised state expanding the potential for regulating the HSR.

**Figure 7:**
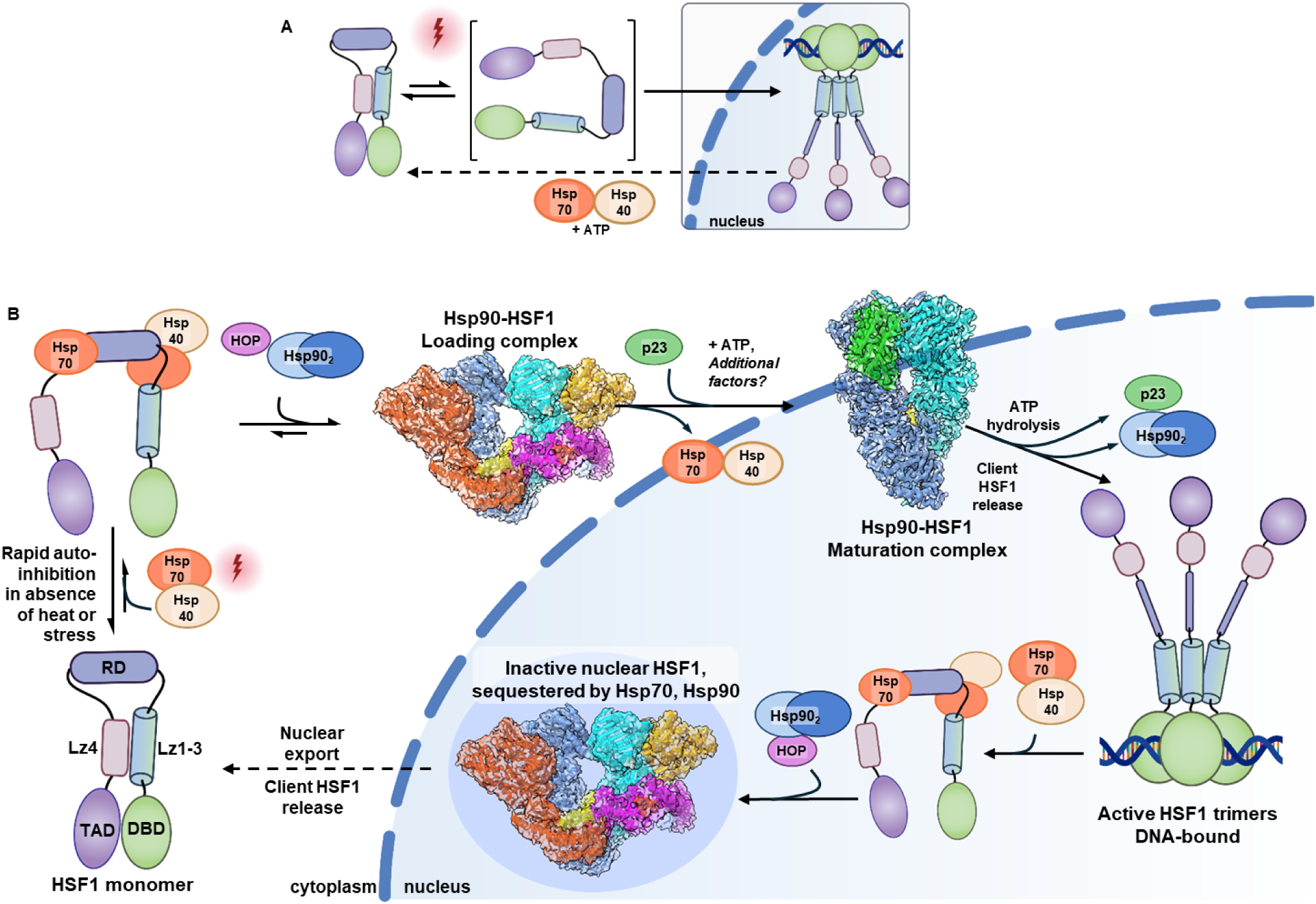
Interlinked Hsp70 and HspG0 activity can both maintain inactive HSF1 and promote HSF1 activation. (A) In a reduced view of HSF1 activity, heat shock drives HSF1 trimerization which promotes nuclear import and binding to target promoters. DnaJB1 (Hsp40) and Hsp70 inactivate HSF1 when stress subsides. (B) Proposed roles of Hsp90 in regulating HSF1 activity. While DnaJB1-Hsp70 are sufficient to disassemble HSF1 trimers, they may also intercept HSF1 monomers in the process of oligomerization. HOP recruits HSF1-DnaJB1-Hsp70 to Hsp90. Whereas loading states may block nuclear import by occluding the HSF1 NLS, closed Hsp90 states promote nuclear import of the client HSF1, especially in conjunction with FKBP52. On closed Hsp90, the Lz1-3 domain of HSF1 is exposed, also promoting activation. As proteotoxic stress subsides, HSPs induced by HSF1 disassemble HSF1 trimers, and Hsp90 complexes either sequester nuclear HSF1 or promote its return to the cytosol. Integration of HSF1 into the Hsp90 cycle may allow for more sensitive stress response, especially to non-heat proteotoxic stresses, and for more efficient nuclear cytoplasmic shuttling to control local HSF1 concentrations.

A major outstanding question regarding Hsp90-HSF1 interactions is how they influence nuclear localization. HSF1 has its own nuclear localization signal (NLS, residues 204-224) (Vujanac, 2005)^35^ that is thought to be occluded in the auto-inhibited state but exposed in the trimer. Our data indicate the NLS is especially exposed in Hsp90 maturation complexes, implying closed states are important for nuclear trafficking (Figure 7B, top). Notably, FKBP51 and FKBP52 bind to closed Hsp90 complexes with opposing roles in nuclear trafficking steroid hormone receptors67, and immunophilin-Hsp90-HSF1 interactions have been reported (Guo, 2001)^46^. Conversely, we also detect DnaJB1-Hsp70 binding at HSF1(202-207), and HOP binding at 241-253 (Figure 4A), which align with DnaJB1 and Hsc70 binding sites reported by Kmiecik et al. (2020)^26^ and an Hsp90 binding site at HSF1(183-214) reported by Kijima et al. 201828 that we did not detect. These Hsp70, HOP, and Hsp90 interactions with HSF1 imply an Hsp90-HSF1 loading complex that likely occludes the NLS and could promote nuclear export. Beyond maintaining a poised intermediate, Hsp90-HSF1 complexes likely play a key role in regulating the cytoplasm:nuclear balance.

Chaperone machinery genes that include *HSPSOAA* and *STIP1* are strongly induced in the HSR (Solis, 2016)^5^, and at higher Hsp90 concentrations, we would expect HSF1 to be driven into Hsp90 complexes. In addition to greater nuclear export, this should promote binding to the secondary Hsp90-HSF1 interaction sites in the regulatory and transactivation domains, blocking recruitment of co-transcription factors by HSF1 (Figures 4A, 7B). Notably, HSF1 is predominantly nuclear in some human cell types (Joutsen, 2024)^68^ and cancers (Mendillo, 2012; Gaglia, 2020)^14,30^, as in yeast (Solís, 2016)^5^, but these cells still have an inducible HSF1-dependent HSR. We expect an Hsp90 loading state to occlude stretches of the RD or TAD to directly block binding of other transcriptional machinery or sequester HSF1 away from transcriptionally active condensates (Dea, 2024)^69^ (Figure 7B, bottom). The IDRs in HSF1 are required for its phase separation (Chowdhary, 2022; Zhang, 2022)^70,71^, and this aspect of Hsp90 interactions with IDR clients is an exciting area for future study.

Our work with HSF1 represents a detailed biochemical investigation of how the Hsp40-Hsp70-Hsp90 chaperone pathway interacts with a client primarily composed of intrinsically disordered regions, analogous to human Tau, TDP-53 and α-synuclein (Lackie, 2017; Rutledge, 2022)^72,73^. Hsp90-Tau interactions have been studied *in vitro* (Karagöz 2014; Oroz 2018; Weickert 2020; Lopez, 2021; Moll 2022)^49,74–77^, and HOP promotes formation of a Hsp70:HOP:Hsp90:Tau complex (Moll 2022)^77^. Surprisingly, these studies report that Hsp90-co-chaperone action on Tau is Hsp40 and nucleotide-independent, in contrast to our findings with HSF1. The prospect of ATP-independent “holdase” chaperoning of certain clients by Hsp70-Hsp90 is very intriguing, but we believe our results also demonstrate the importance of testing complete systems with active chaperone ATPase activity. Nonetheless, the findings that Hsp90:Tau promotes formation of short oligomers while suppressing aggregation should be analogized to our findings on Hsp90:HSF1, and further highlights that the role of Hsp90 in HSF1 phase separation needs addressing.

Despite the differences between our Hsp90-HSF1 structures and those of other available Hsp90:client complexes, we do not see this as pointing to a distinct Hsp90 mechanism. Unstructured client is regularly present in the Hsp90 lumen, induced by Hsp70 or a co-chaperone like Cdc^37^. Other differences in bound clients’ architecture are likely due to the particulars of the clients or specific client-co-chaperone interactions rather than the requirements for Hsp90 binding. In human cells, open Hsp90 states appear to be much more populated than closed states (Southworth, 2008; Lopez, 2021; Jussupow, 2022; Reidl 2024)^76,78–80^. Although in closed states the client is temporarily locked in, many Hsp90 clients associate transiently (Kolhe, 2023; Taipale, 2012)^31,81^, and we expect more dynamic binding of HSF1 to Hsp90 in a loading state to be physiologically relevant. We found that both elevated temperatures and ATP-competitive Hsp90 inhibitors biased Hsp90-HSF1 complexes towards the loading state, which is then consistent with HSF1 activation by heat stress and in response to Hsp90-targeted therapeutics.

In this work, we use site-directed unnatural amino substitutions to track specific protein-protein interactions in complex biochemical mixtures and as a powerful enabling tool in cryo-EM structure determination. Whereas previous work required mutations that stabilize the closed Hsp90 state to observe appreciable complexes, we observed continuous Hsp90-HSF1 interactions throughout the loading and maturation complexes with active Hsp90. This precise crosslinking greatly facilitated determining cryoEM structures of these complexes, eliminating the need for the general glutaraldehyde stabilization that was previously required and that negatively impacted sample heterogeneity. On the one hand, use of BPA crosslinking has clear utility for studying other client chaperone systems, especially the wide diversity of Hsp90 clients whose chaperoning has not been investigated biochemically. On the other hand, the outstanding questions regarding HSF1 regulation, as described above, will require *in vivo* experimentation. In yeast, Hsp90-client photocrosslinking yielded very interesting results (Kolhe, 2023)^31^. This can be extended to human cells (Baischew, 2023)^82^, where photocrosslinking has the unique advantage of providing a snapshot of direct interactions.

## Acknowledgements

This work was supported in part by NIH grant R35GM118099 (D.A.A.). T.W.O was a Damon Runyon Fellow supported by the Damon Runyon Cancer Research Foundation DRG #2407-20. T.M.P.C. is supported by the National Cancer Institute of the National Institutes of Health (F32CA298768). Cryo-EM equipment at UCSF is partially supported by NIH grants S10OD020054, S10OD021741 and S10OD026881 and Howard Hughes Medical Institute.

## Supplemental Figures

**Figure S1:**
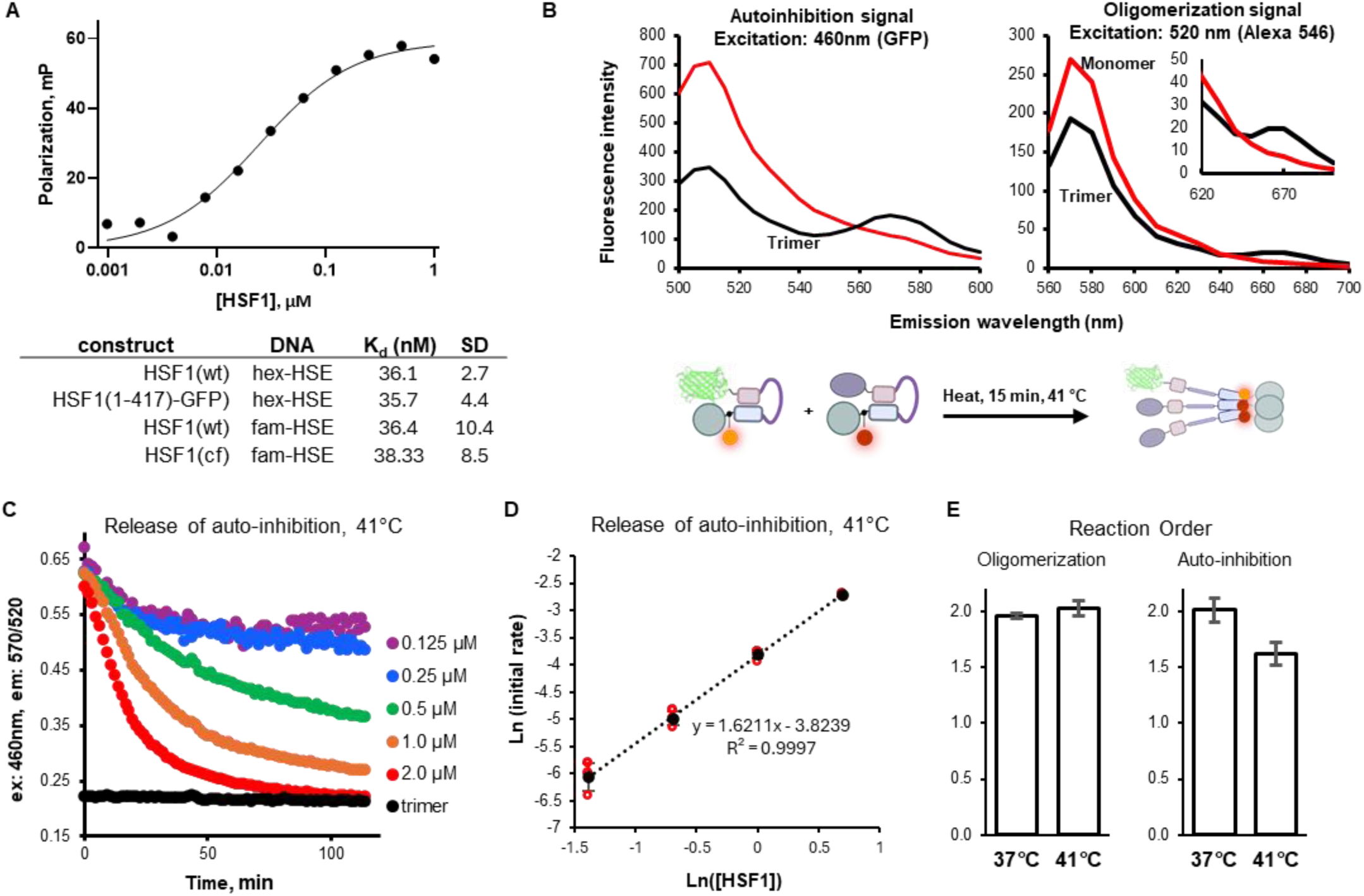
HSF1(fret) system monitors HSF1 auto-inhibition and trimerization. (A) Representative binding curve of HSF1(wt) trimers to 5’ FAM-labeled dsDNA oligos containing the canonical HSE. HSF1 concentrations shown are monomer equivalents. Below, Kd values for trimeric HSF1 binding to HSE oligos labeled with 5’FAM or 5’HEX, n=3-5. (B) Left: Emission scan of monomeric and trimeric HSF1(cf, 1-417, L125C-Alex546)-GFP when excited at 460 nm. Right: Emission of scan of HSF1(fret), equimolar mixture of HSF1(cf, 1-417, L125C-Alex546)-GFP and HSF1(cf, L125C-Alex647), excited at 520 nm. Below: Scheme of conformational changes during trimerization. (C) Monomer concentration affects the rate of HSF1(fret) autoinhibition signal decrease during heat shock. Representative curves. (D) Initial rates of auto-inhibition release were calculated for varying HSF1(fret) concentrations and plotted Ln(rate) vs Ln([HSF1]). Experiments were performed in triplicate. The slope of the linear fit line is the order of reaction with respect to HSF1 concentration. (E) Order of reaction was calculated for auto-inhibition release and oligomerization at 37 °C and 41 °C. Error bars denote standard deviation of three experiments.

**Figure S2:**
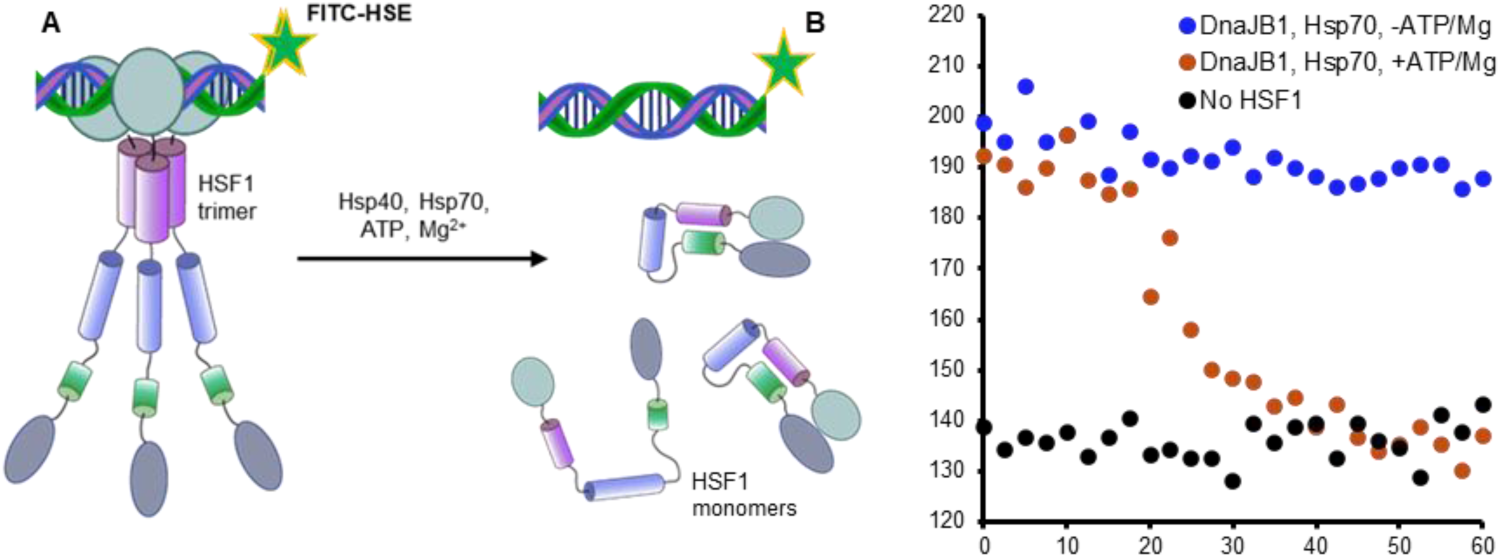
DnaJB1, Hsp70, and ATP-Mg displace HSF1 trimers from HSEs. (A) Monomerization of HSF1 trimers releases bound 5’FAM-HSE. (B) HSF1, FAM-HSE, and chaperone components were co-incubated at 28 °C. At the start of the time series, addition of 3 mM ATP + 4 mM MgCl2 to a mixture of 0.8 µM trimeric HSF1 (2.4 µM monomer equivalent), 100 nM 5’FAM-HSE, 5 µM DnaJB1 (monomer concentration), and 10 µM Hsp70 leads to decrease in fluorescence polarization indicative of HSF1 monomerization.

**Figure S3:**
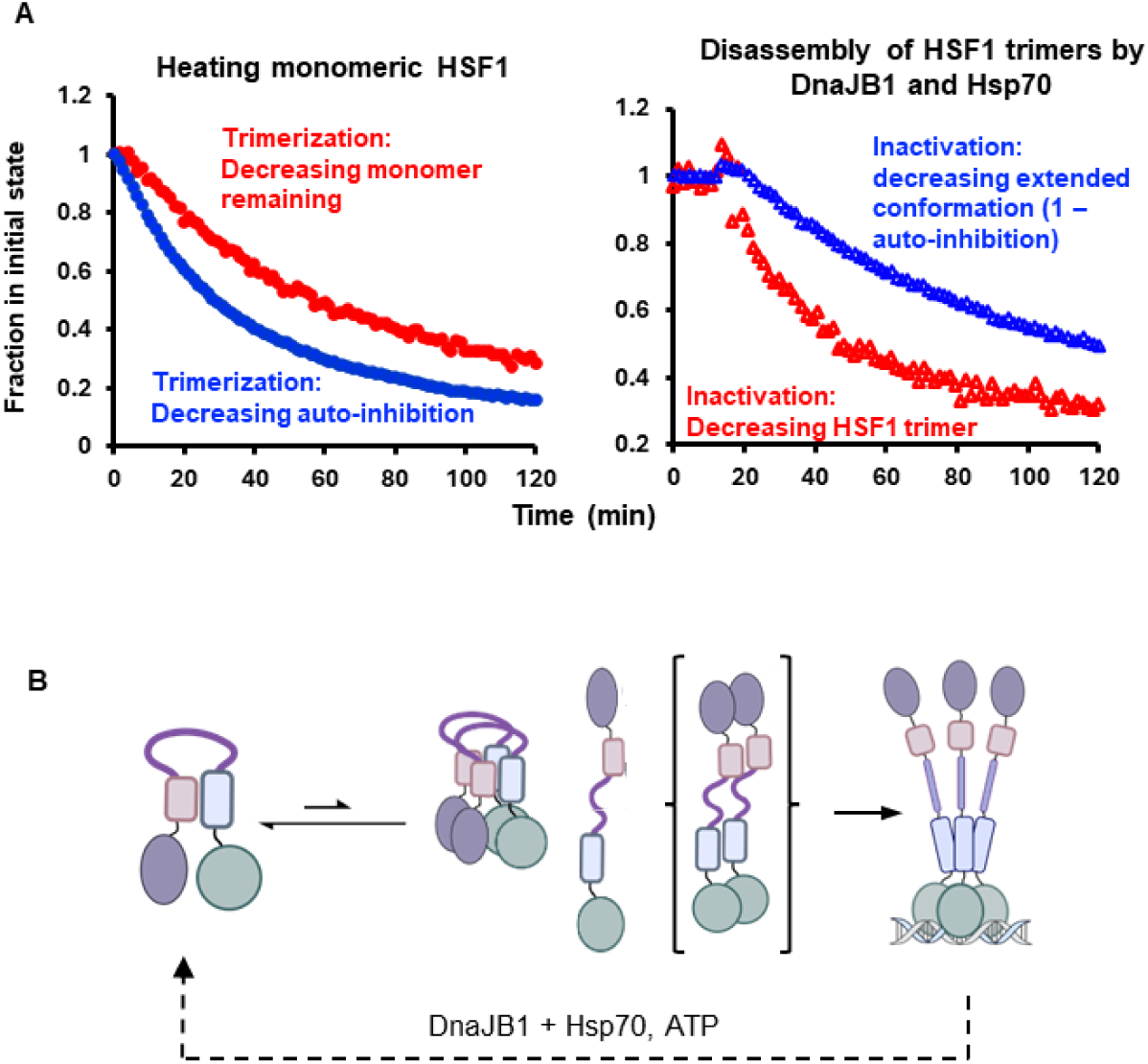
Relative rates of change in oligomerization and auto-inhibition. (A) Auto-inhibition and oligomerization FRET signals from heating and monomerization experiments were normalized to the signal of HSF1 monomers and trimers. In heating experiments, the auto-inhibition signal decreases faster than oligomerization signal increases (displayed as decreasing monomer). When monomerizing HSF1(fret) trimers with DnaJB1-HSp70, the oligomerization signal decreases faster than the auto-inhibition signal increases (displayed as decreasing trimer, or 1 – scaled auto-ihibition). (B) Possible intermediate states of HSF1 include auto-inhibited dimers, extended monomers, and a short-lived extended dimer.

**Figure S4:**
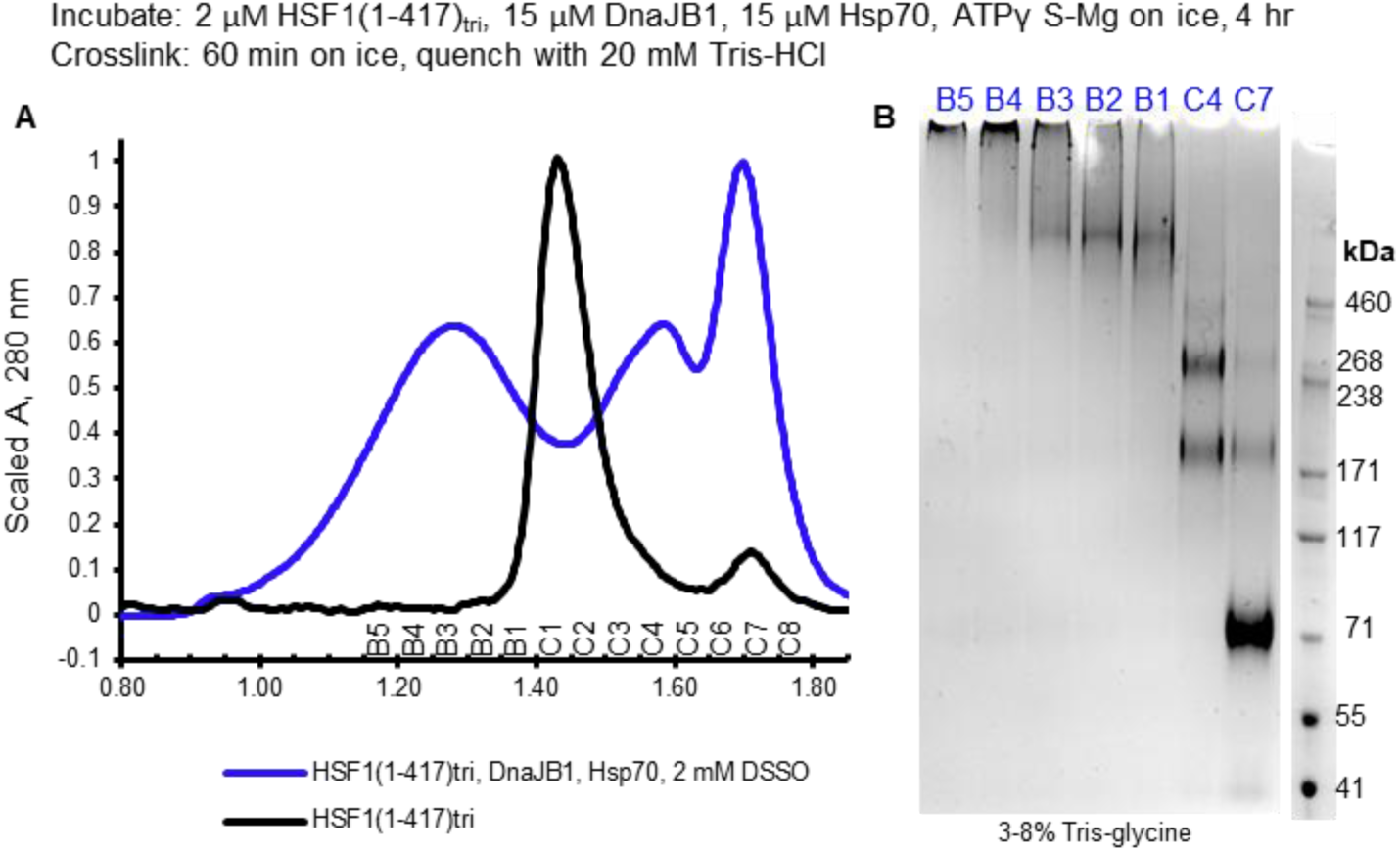
DnaJB1, Hsp70, HSF1 form a high molecular weight complex in the presence of a non-hydrolyzable ATP analog. (A) Analytical size exclusion chromatography traces of HSF1(1-417) trimers and HSF1(1-417)-DnaJB1-Hsp70 complexes DSSO-crosslinked in the presence of ATPγ S-Mg. (B) SDS page analysis of fractions from (A) shows a show a single, high molecular weight band in the SEC peak.

**Figure S5:**
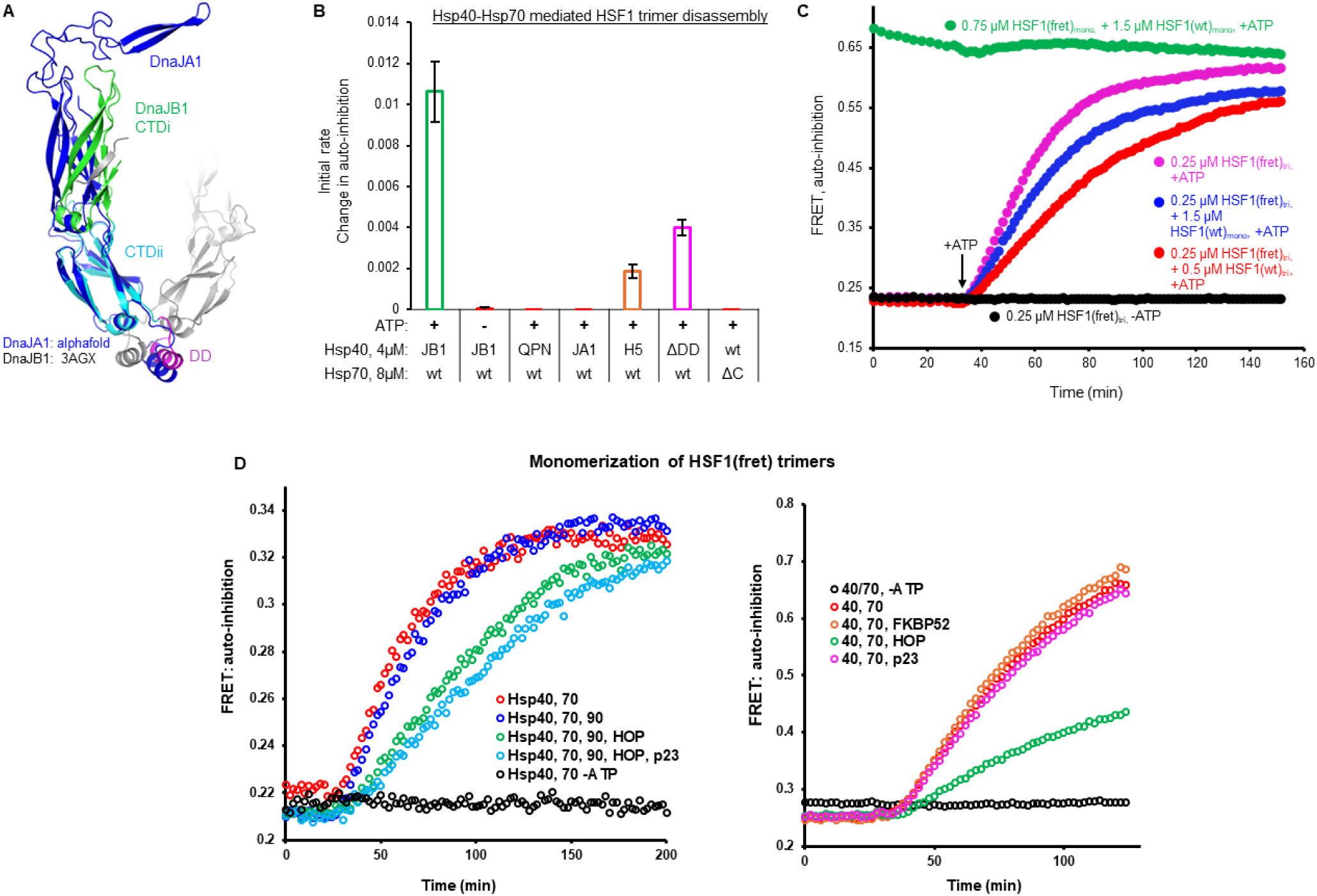
Determinants of HSF1 monomerization by Hsp40-Hsp70. (A) Alignment of the C-terminal domains (CTDs) and dimerization domains (DDs) of DnaJA1 (Alphafold #AF-P31689-F1) and a crystal structure of DnaJb1 (#3AGX). (B) DnaJB1 with Hsp70, but not DnaJA1, is able to monomerize HSF1(fret) trimers. Mutations which block DnaJBA1 J-domain function (QPN), auto-inhibition (H5), and dimerization (ΔDD), block or greatly reduce the rate of HSF1 monomerization. Removal of the EEVD motif from the C-terminus of Hsp70 completely abrogates its activity with DnaJB1. (C) Doping in unlabeled HSF1(wt) trimers slows monomerization of HSF1(fret) trimers to a greater extent than addition of HSF1(wt) monomers. (D) Effects of Hsp90 and co-chaperones on the rate of auto-inhibition increase during DnaJB1-Hsp70-mediated monomerization of HSF1(fret) trimers

**Figure S6:**
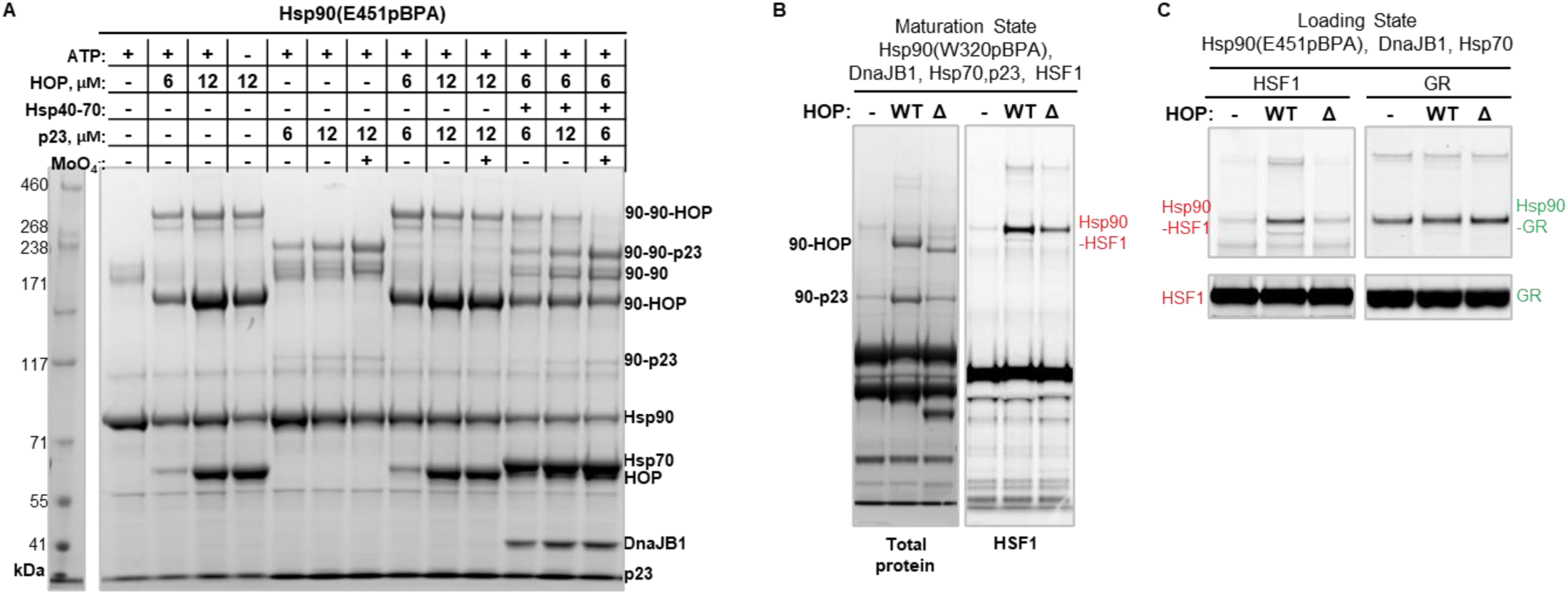
HOP inhibits Hsp90 closure and positions HSF1 in the HspG0 loading state. (A) HOP blocks p23 induced closure of p23, but this is relieved by Hsp70. Where indicated, components were included at the following concentrations: 12 µM Hsp90 (monomer), 12 µM Hsp70, 4 µM DnaJB1, 3 mM ATP, 10 mM NaMoO4. HOP and p23 were used at the concentrations shown. Samples were incubated 90 minutes at 28 °C. (B) HOP strongly promotes HSF1 interaction with Hsp90(W320pBPA) in the closed state, and deletion of the HOP DP2 domain (Δ) reduces HSF1-Hsp90 crosslinking. These experiments used 20 µM Hsp90, 20 µM Hsp70, 6 µM DnaJB1, 10 µM HOP or HOP(ΔDP2), 20 µM p23, and 1 µM HSF1(cf, Y225C-Alexa647), incubated 90 minutes at 28 °C. (C) HOP strongly promotes HSF1 interaction with Hsp90(E451pBPA) in the loading state, and DP2 domain deletion reduces HSF1-Hsp90 crosslinking. In contrast, GR is much less dependent on HOP under these conditions, and DP2 domain deletion has little effect. These experiments used 20 µM Hsp90, 15 µM Hsp70, 4 µM DnaJB1, 8 µM HOP, and 1 µM MBP-GRLBD (labeled with sub-stoichiometric Alexa 488 TFP) or HSF1(cf, Y225C-Alexa647).

**Figure S7:**
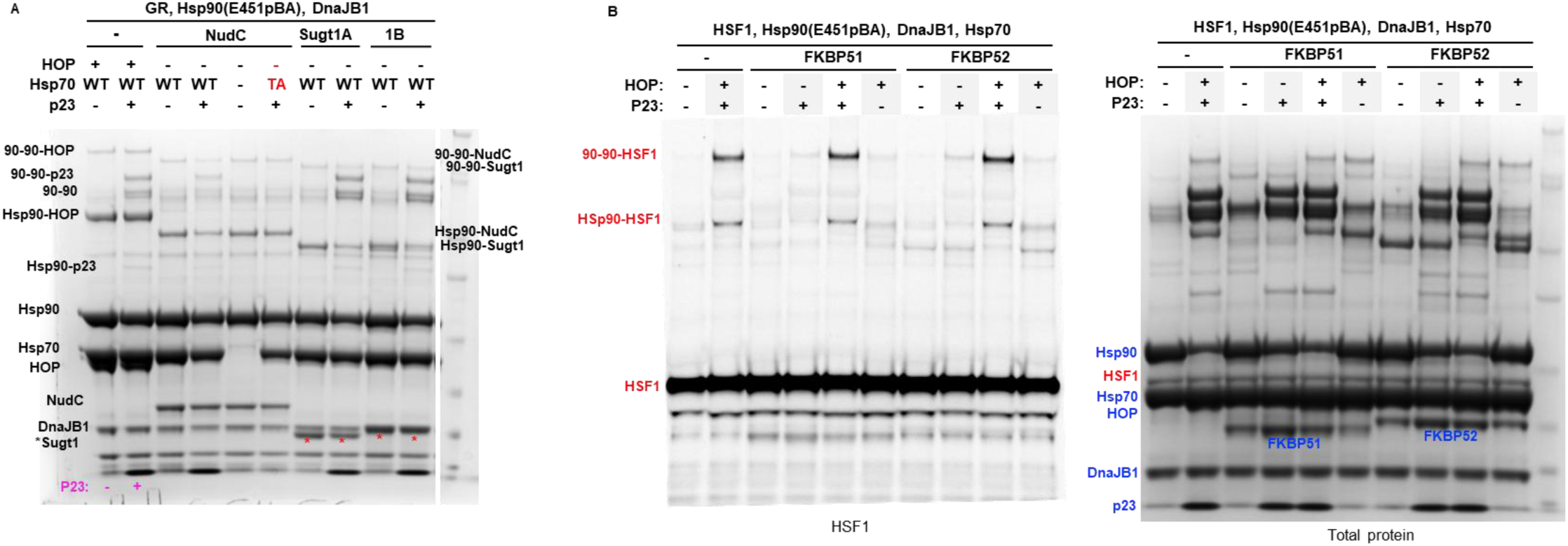
Effects of other co-chaperones on Hsp90-client crosslinking systems. (A) NudC and Sugt1 isoforms, like HOP, interact with the Hsp90 middle domain at E451pBPA. NudC inhibits p23 induces Hsp90 closure but Sugt1 does not. (B) In-gel HSF1(cf, Y225C-alexa647) fluorescence shows immunophilins FKBP51 and FKBP2 are compatible with HSF1-Hsp90 complexes but do not have a significant effect on the strength of the interaction. Total protein staining shows FKBP52, but not FKBP51, crosslinks at HSp90(E451pBPA) in both (apparently) open and closed Hsp90 states.

**Figure S8:**
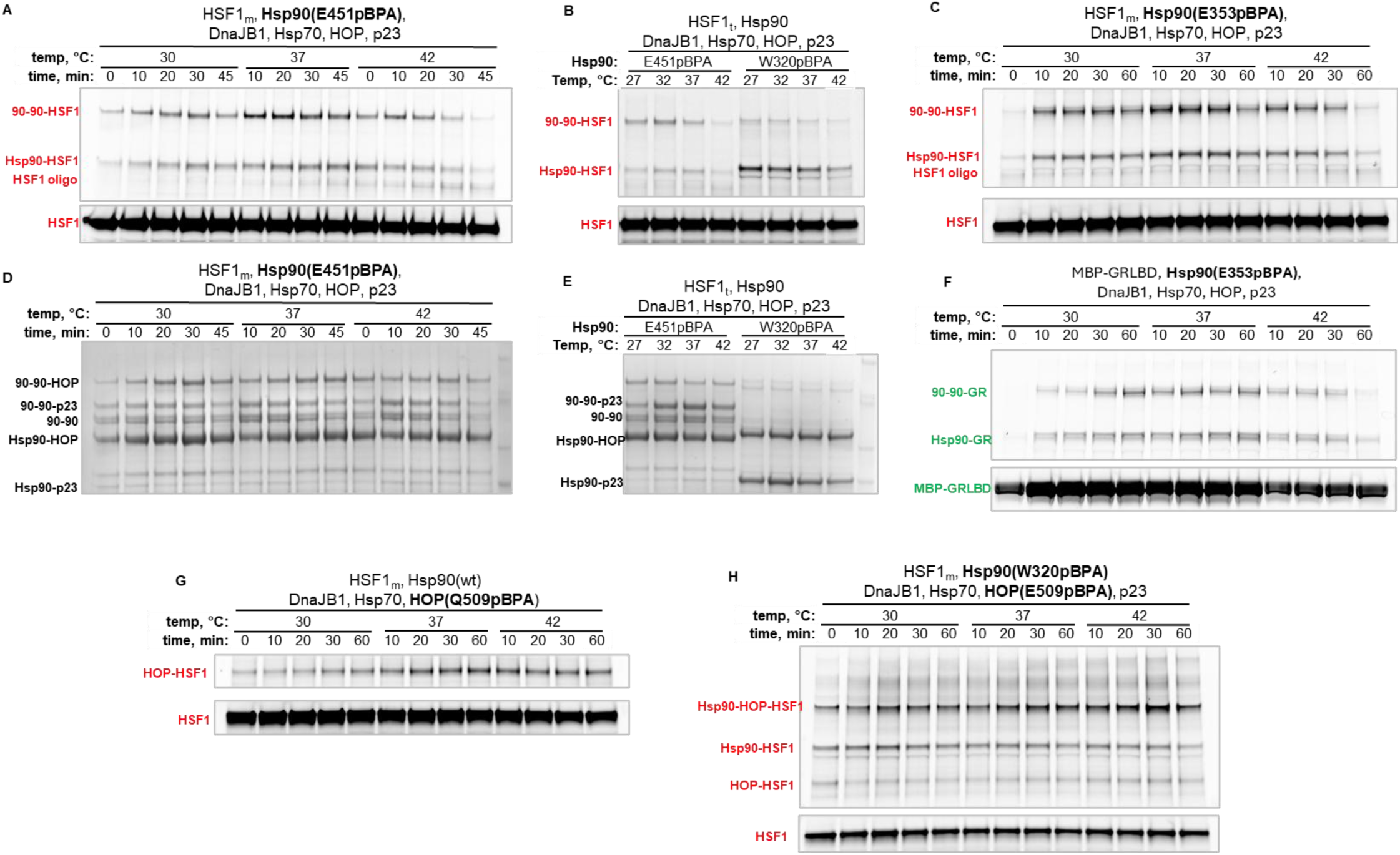
Temperature modulates Hsp90 conformation and HSF1-HspG0 interactions. For these experiments, reactions containing 20 µM Hsp90(wt or XpBPA), 15 µM Hsp70, 10 µM HOP, 4 µM DnaJB1, 20 µM p23, and 2 µM HSF1(cf, Y225C-Alexa 647) monomers were assembled at room temperature, started by the addition of 3 mM ATP, then moved to the indicated temperatures. Differences in components or concentrations are noted below. (A) Time course of HSF1-Hsp90(E451pBPA) interactions show temperature-dependent dynamics; at higher temperatures the maturation state becomes more rapidly populated, then decays. At 37 °C, buildup of loading state HSF1-Hsp90 is apparent at later timepoints, whereas at 42 °C increased HSF1 oligomer is seen. (B) Temperature series comparing Hsp90(E451pBPA) and Hsp90(W320pBPA) shows decreased HSF1 crosslinking at higher temperatures. (C) Time course experiment similar to (A), using 20 µM Hsp90(E353pBPA). The effect of heat on the crosslinking pattern does not change with pBPA substituted at a different site. (D) Total protein staining of the gel shown in (A); Hsp90-Hsp90-p23 crosslinking peaks faster and decreases sooner at higher temperature. (E) Total protein staining of the gel in (B). (F) A parallel experiment to that shown in (C) was run using 2 µM MBP-GRLBD (Alexa Fluor 488 labeled) rather than HSF1; a similar effect is seen with heating to 42 °C. (G) Higher temperatures favor HSF1-HOP(E509pBPA) crosslinking in a reconstitution with loading state components, and crosslink strength does not decrease at longer timepoints as seen in (A) and (C). For this experiment, p23 was excluded and 10 µM HOP(E509pBPA) was used in place of HOP(wt). (H) In the loading state, Hsp90(W320pBPA) crosslinks HOP strongly and HSF1 very weakly in contrast to closed Hsp90(W320pBPA), which crosslinks HSF1. Including both Hsp90(W320pBPA) and HOP(E509pBPA) thus allows us to assess both states with higher confidence in the same experiment; 10 µM HOP(E509pBPA) was used in place of HOP(wt).

**Figure S9:**
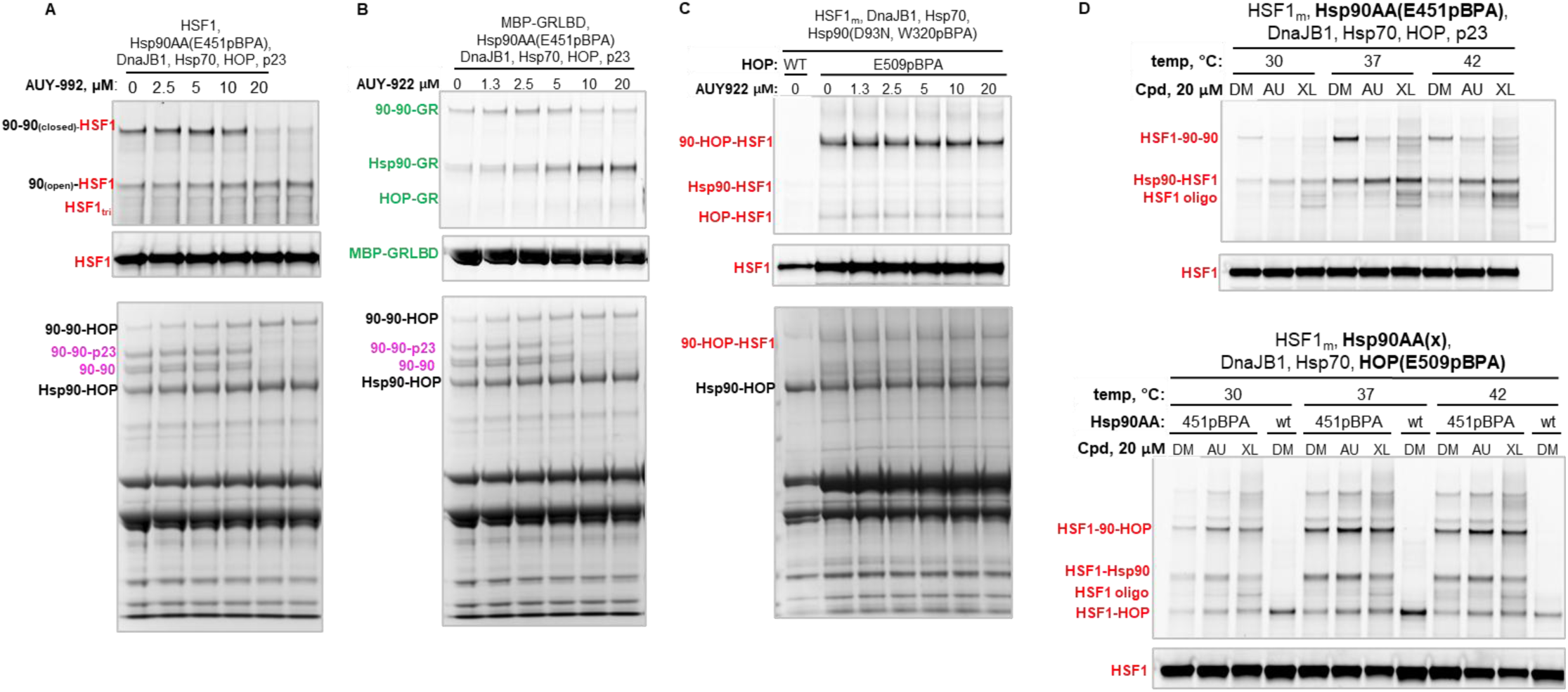
Effects of ATP-competitive Hsp90 inhibitors on HSF1-HspG0 interactions. For these experiments, reactions containing 20 µM Hsp90, 15 µM Hsp70, 10 µM HOP, 4 µM DnaJB1, 20 µM p23, and 2 µM HSF1(cf, Y225C-Alexa 647) monomers were assembled at room temperature, started by the addition of 3 mM ATP, then moved to the indicated temperatures. Differences in components or concentrations are noted below. (A) HSF1 and chaperone components were mixed as a single reaction, then aliquoted and supplemented with DMSO (vehicle) or AUY-922 to the same final DMSO concentration. Reactions run 75 minutes at 28 °C. Top panels show in gel Alexa Fluor 647 fluorescence; bottom panel shows total protein staining. (B) Experiment run in parallel with (A), using 1 µM MBP-GRLBD in place of HSF1. Top panels show in gel Alexa Fluor 488 fluorescence; bottom panel shows total protein staining. (C) 20 µM Hsp90(D93N, W320pBPA) and 10 µM HOP(E509pBPA) were used to assess HSF1 bound in the loading state. Conditions were otherwise the same as (A) (D) Top panel: Reactions were run at 28 °C for 45 minutes, then moved to the indicated temperature for 15 minutes. DMSO (DM), AUY-922 (AU), or XL-888 (XL) were added to the reactions to equal DMSO concentration prior to the addition of ATP. Samples were moved to ice and supplemented with 10 mM NaMO4 at the end of heating, prior to crosslinking. Bottom panel: Setup as in the top panel, except that either 20 µM Hsp90 wt or E451pBPA was used as indicated, with 10 µM HOP(E509pBPA) to capture loading states. In place of NaMO4, 3 mM ADP was added to stabilize the loading state.

**Figure S10:**
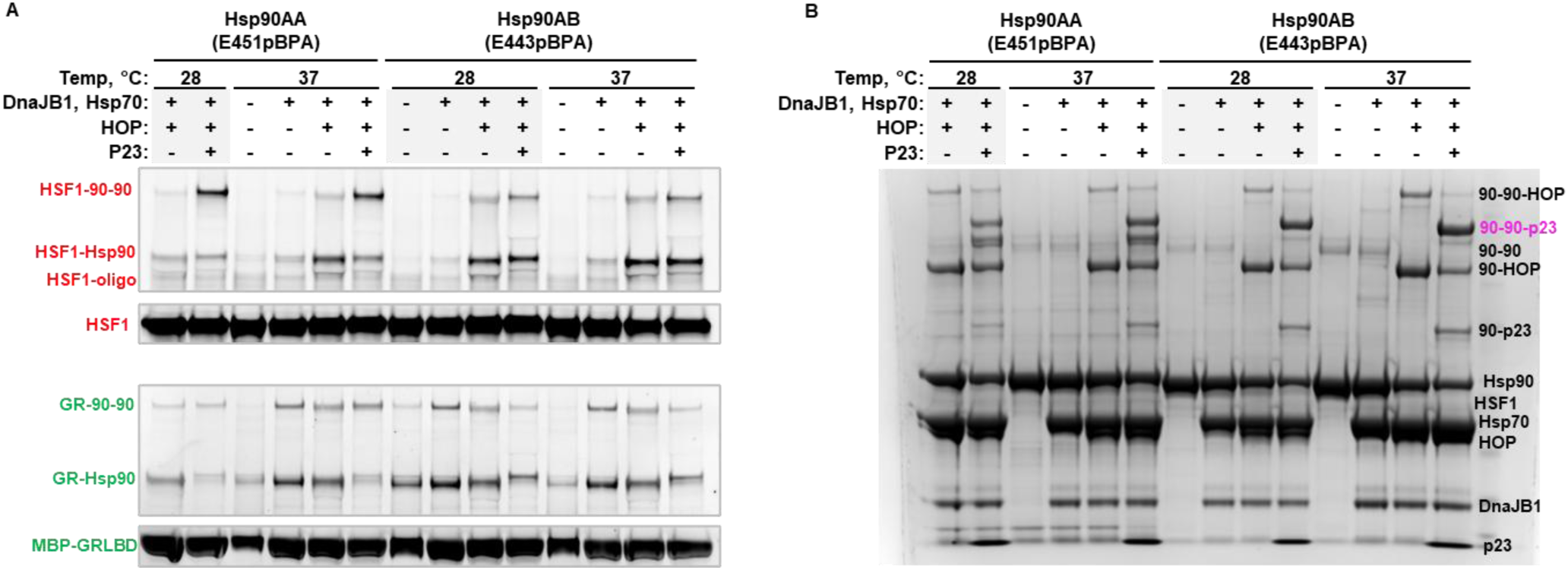
HspG0AA and HspG0AB have similar interactions with HSF1, as well as co-chaperones. Indicated components were used at the following concentrations: 20 µM Hsp90, 15 µM Hsp70, 10 µM HOP, 4 µM DnaJB1, 20 µM p23, and 1.5 µM HSF1(cf, Y225C-Alexa 647) monomers or 1.0 µM MBP-GRLBP. Reactions were started by the addition of 3 mM ATP, then moved to 28 °C or 37°C for 1 hr. Then, 10 mM NaMO4 was added to reactions containing p23, all reactions incubate additional 30 minutes prior to UV exposure at 4 °C. (A) In-gel fluorescence of HSF1 (top panels) and GR (bottom panels) (B) Total protein staining of the HSF1 gel in panel (A). Note that for Hsp90AB(E443pBPA) the Hsp90-Hsp90-p23 crosslinking is stronger than seen with Hsp90AA(E451pBPA). This may explain the relative weakness of the Hsp90-Hsp90-HSF1 crosslinks in the Hsp90AB samples; that is, differences in pBPA position or orientation may favor p23 crosslinking over HSF1 crosslinking in the Hsp90AB maturation state.

**Figure S11:**
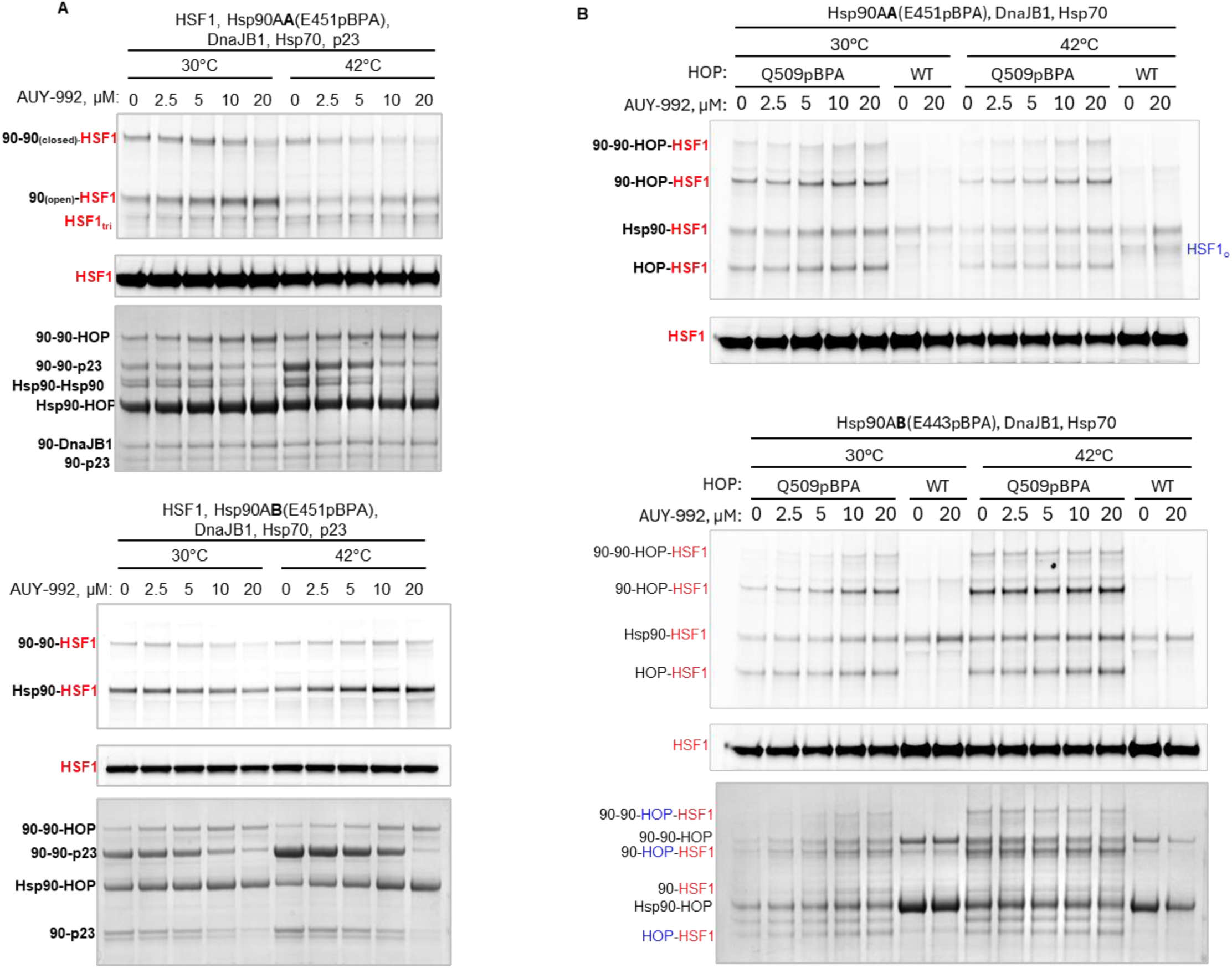
ATP-competitive inhibitors differentially affect HSF1 interactions with HspG0AA and HspG0AB. Indicated components were used at the following concentrations: 20 µM Hsp90, 15 µM Hsp70, 10 µM HOP, 4 µM DnaJB1, 20 µM p23, and 3 µM HSF1(cf, Y225C-Alexa 647). Reactions were started by the addition of 3 mM ATP, then moved to 28 °C for 1 hr. Then, samples were heated to 30 °C or 42 °C as indicated for 15 minutes. Next, 10 mM NaMO4 was added to reactions containing p23 and 3 mM ADP added to the others for 3 minutes before samples were moved to ice (5 minutes) and exposed to UV at 4 °C. (A) In-gel fluorescence of HSF1 (top panels) and total protein staining (bottom panels). Hsp90-Hsp90-p23 crosslinking trends are the same for the two HSp90 isoforms, but Hsp90-HSF1 interactions are affected differently by AUY-922. Note that 90AA-HSF1 crosslinks increase with more inhibitor at 30 °C and 42 °C; in contrast, 90AB-HSF1 crosslinks weaken with increasing inhibitor at 30 °C. (B) To assess HSF1 in Hsp90 loading states, HOP(E509pBPA) was used in place of HOP(wt). Bottom panel shows total protein staining for the experiment with HSp90AB(E443pBPA).

**Figure S12:**
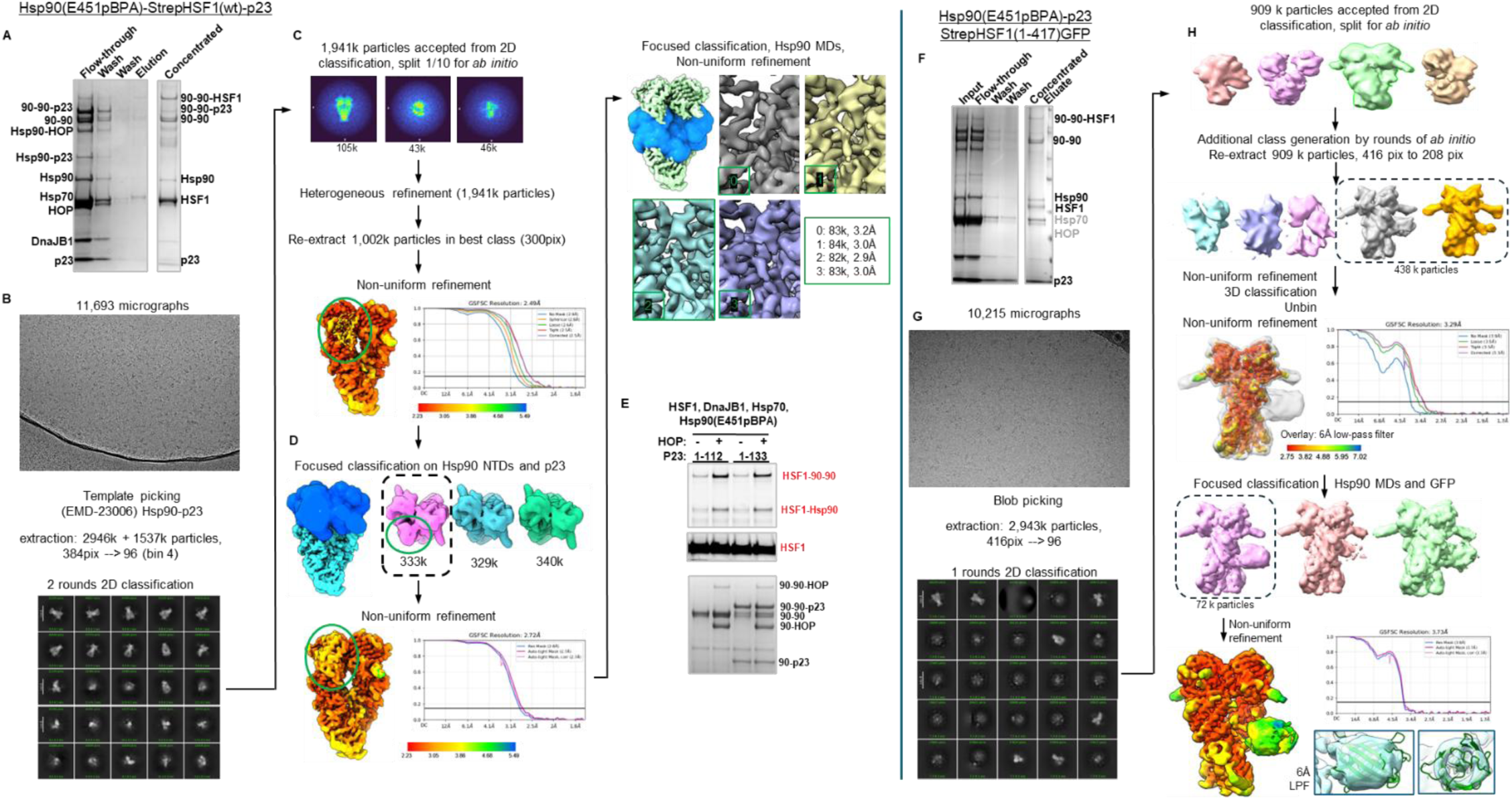
Purification of closed state Hsp90-HSF1 complexes and single particle cryo-EM processing pipeline. A: Coomassie-stained SDS-PAGE gel showing purification of photo-crosslinked Hsp90(E451pBPA)-HSF1-p23 complexes via Strep-Tactin affinity resin. B: A representative micrograph shows relatively uniform particles without additional samples purification steps. Template picking and 2D classification were used to select particles for 3D reconstruction. C: Ab initio reconstruction with a subset of particles was used to generate initial 3D classes, particles were sorted by heterogeneous refinement, and those in the one good class were re-extracted and aligned to an initial consensus map. D: Density for p23 was weak relative to Hsp90 in the consensus map and focused 3D classification was performed to select particles with p23 present. The mask used is shown in blue at left. In the refined map of the Hsp90(E451pBPA)-p23-HSF1 map (bottom), the crosslinks seen are predominantly Hsp90-Hsp90, but other modes are likely present. An addition round of focused 3D classification on the HSp90 MDs and HSF1 revealed heterogeneity but did not improve interpretability. E: SDS-PAGE followed by in-gel fluorescence and Coomassie staining show that full deletion of the p23 tail, p23(1-112), eliminates Hsp90(E451pBPA)-p23 crosslinking but does not block Hsp90 closure (90-90 crosslinks preserved) or visibly alter Hsp90-HSF1 interactions. F: Coomassie-stained SDS-PAGE gel showing purification of photo-crosslinked Hsp90(E451pBPA)-HSF1(1-417)GFP complexes via Strep-Tactin affinity resin. Not all Hsp70 and HOP were washed out in this sample but did not appear in subsequent processing steps. G: A representative micrograph shows somewhat heterogeneous particles uniform particles, and blob picking and 2D classification were used to select particles for 3D reconstruction. H: Several rounds of ab initio reconstruction generated two classes with clear density for Hsp90, HSF1, and additional density. One contained p23 as well (grey in dashed box). A consensus map was refined with these particles, and used to generate additional 3D classes; notably, p23 and GFP were not present in the same classes although luminal HSF1 density was present in either case. The GFP-containing classes were refined together, and then focused classification was performed on the Hsp90 MDs and GFP. The final map contains a clear barrel-shaped region that well accommodates a GFP (bottom, inset).

**Figure S13:**
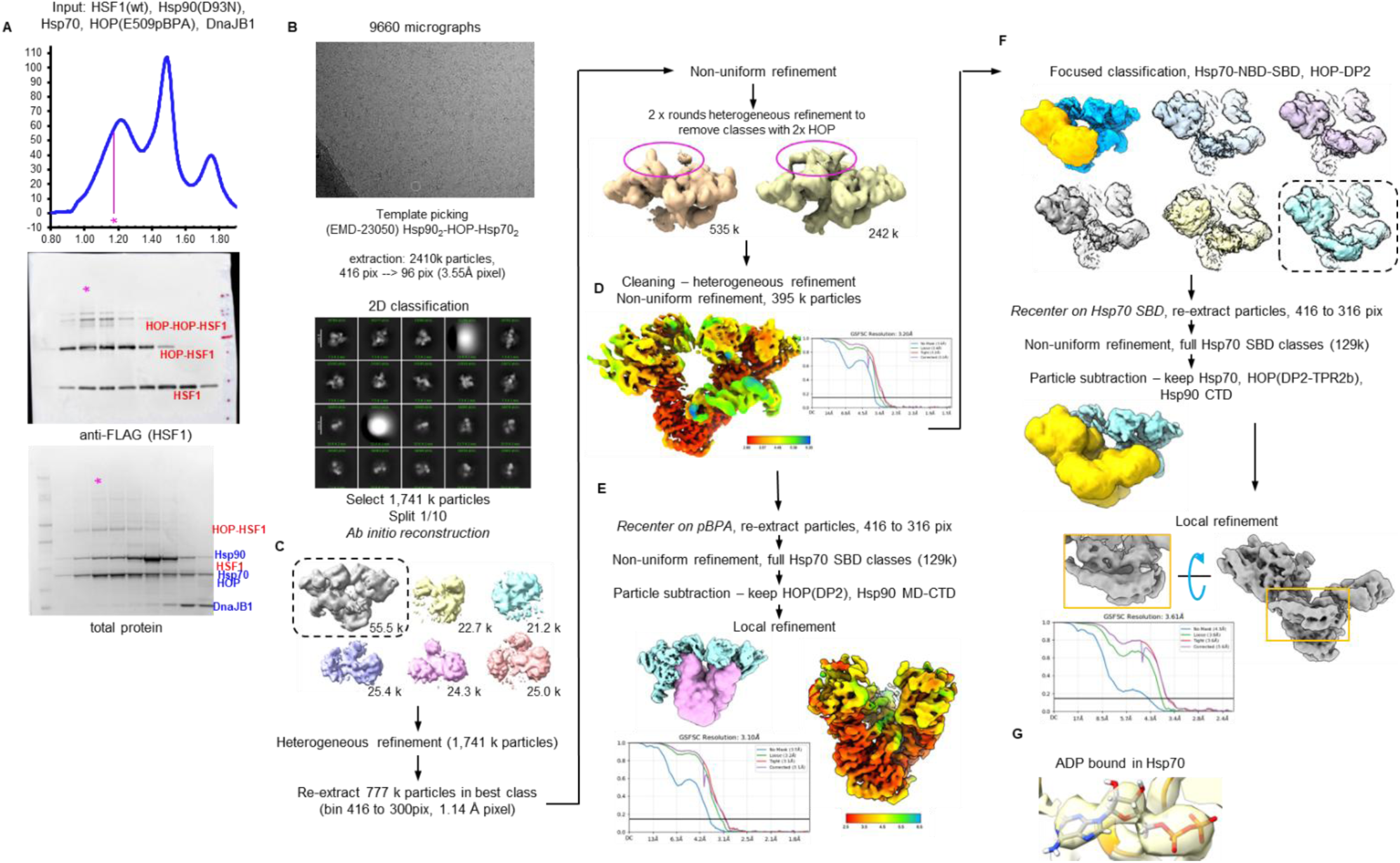
Purification of loading state Hsp90-HSF1 complexes and single particle cryo-EM data processing workflow. A: A cycling *in vitro* reconstitution of HSF1(wt), Hsp90(D93N), Hsp70, HOP(E509pBPA), and DnaJB1 was photo-crosslinked and then directly subjected to size-exclusion chromatography without additional purification of stabilization steps. Collected fractions were analyzed by SDS-PAGE then western blot and Coomassie staining to identify which fractions contained both HSF1-HOP(E509pBPA) adducts and other chaperone components. The fraction marked with a star was used for cryo-EM analysis. B: Particles were picked from selected micrographs using templates generated from the Hsp902-Hsp702-HOP-GR structure (EMDB: 23050) with GR removed. 2D classification was used to curate particles. C: Ab initio reconstruction of selected particles generated one class that strongly resembled an Hsp90 loading state, and heterogeneous refinement was used to sort between that class and junk classes. The consensus class had weak density for a second copy of HOP (as seen by western blot, A). Differential filtering and thresholding were used to make two classes, with one or two copies of HOP, and two rounds of heterogenous refinement were run to remove particles with two copies of HOP (center, top, highlighted in pink). D: The cleaned consensus reconstruction resolves two copies of Hsp90, and two Hsp70 NBDs, HOP domains TPR2a-TPR2B-DP2, however, the Hsp90 CTDs and MDs are much clearer than other regions. E: To better resolve the client HSF1, particles were recentered on HOP(E509pBPA), the consensus map refined, then particle subtraction and local classification performed targeting the Hsp90 CTDs, MDs, and HOP DP2 domain F: Multiple states of the Hsp70 SBDs were observed. Focused classification on the Hsp70 (NBD and SBD) near HOP showed multiple conformations of the SBD; in one of these both the beta domain and the alpha helical lid were resolved. Local refinement with particle subtraction better resolved The HSp70 SBD and bound HSF1 client. G: ADP bound in the locally refined Hsp70 SBD.

**Figure S14:**
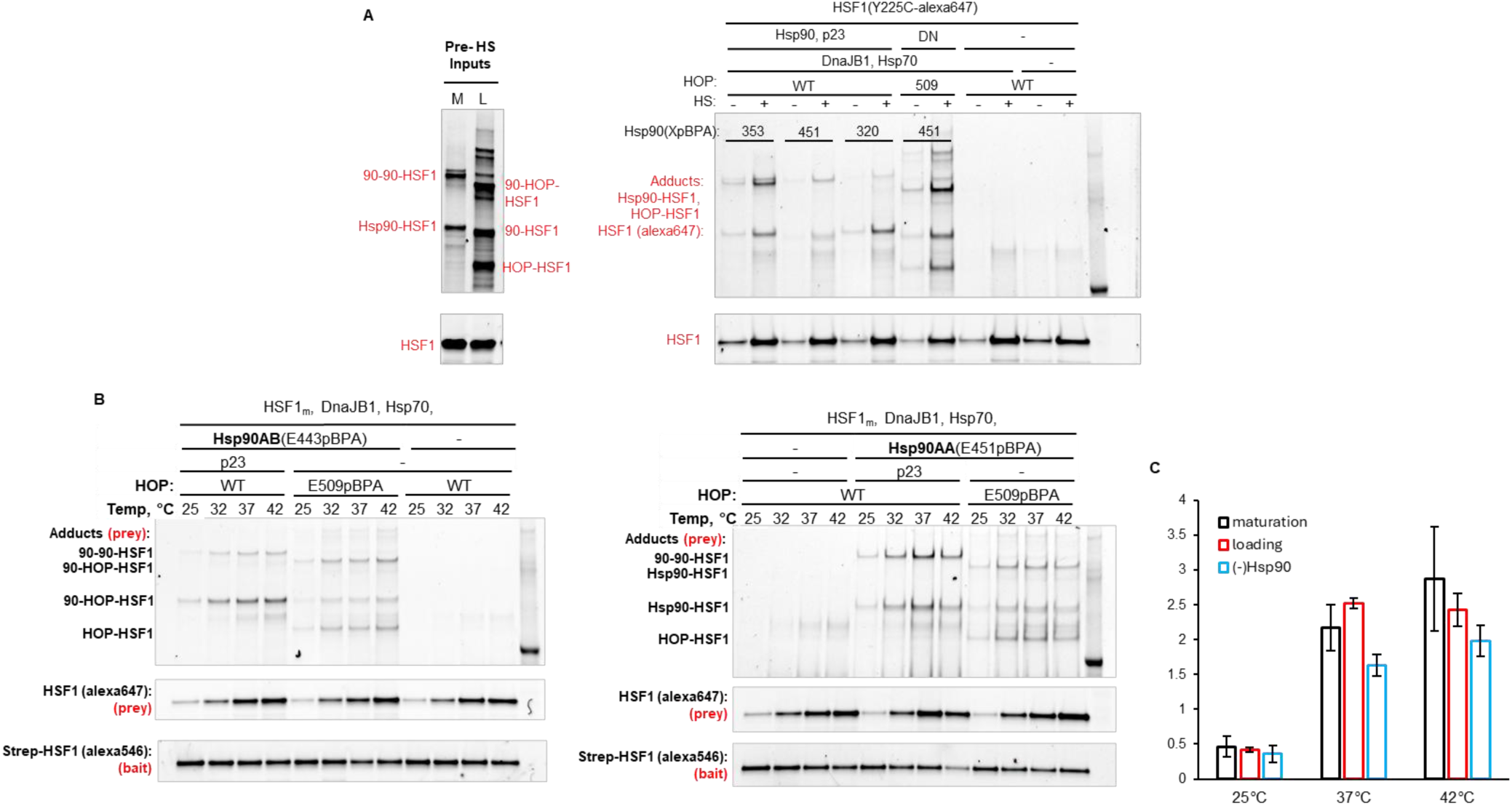
Results of HSF1 Bait-Prey experiments are not sensitive to pBPA position or HspG0 isoform. (A) Samples comprising 3 µM HSF1(cf, Y225C-alexa647) and chaperones were incubated 1 hr, 28 °C. As indicated, 20 µM Hsp90, 20 µM Hsp70, 10 µM HOP, 4 µM DnaJB1 and 20 µM p23 were used. For the maturation state, p23 added after 30 minutes, and NaMoO4 after 50 minutes. Samples were moved to ice, then exposed to UV for 1 hr at 4 °C. The left panel shows the results of this crosslinking reaction for example maturation (M) and loading (L) state samples, the inputs to the pulldown. The right panel shows the adducts and free HSF1 pulled down by bait twinStrep-HSF1(wt) with series of chaperone combinations, with and without heat shock prior to Strep-Tactin pull-down. (B) Side-by-side bait-prey experiments were performed with either Hsp90AA(E451pBPA) and Hsp90AB(E443pBPA). Overall, both HSp90 isoforms form complexes with HSF1 in which HSF1 can be trimerized with free HSF1.

## Methods

### Purification and labeling of HSF1 and GR constructs

*E. coli* codon optimized human HSF1 was cloned into a pCDFduet vector with Ulp1 catalytic domain (ulp1cat, residues 403-621) in first site and SUMO-FLAG-3C-*hs*HSF1-his6 in the second site. This vector was transformed into BL21(DE3) *E. coli* for heterologous expression. Overnight cultures were diluted 1:100 into fresh terrific broth + 100 µg/mL spectinomycin + 1.0 mM MgCl2, and grown at 37 °C to OD600 ∼ 1.0, cooled to 18 °C, and induced and then induced with 200 µM Isopropyl β-D-1-thiogalactopyranoside (IPTG, Fisher Scientific)). After overnight expression, cells were harvested, resuspended in lysis buffer (500 mM NaCl, 30 mM Tris-HCl pH 8.0, 10% v/v glycerol), and supplemented with 10 µg/mL lysozyme (Sigma Aldrich),), 1.0 µg/mL bovine DNaseI (GoldBio),1.0 mM phenylmethylsulfonate fluoride (PMSF, Thermo Fischer), cOmplete protease inhibitor (Roche), 3 mM β-mercaptoethanol (βMe), 0.1% TWEEN-20, and 1 mM ethylenediaminetetraacetic acid-NaOH pH 8.0 (EDTA). Resuspended cells were homogenized with an immersion blender and lysed by 3x freeze-thaw in liquid nitrogen. Following the last thaw, lysates were clarified by ultracentrifugation at 100,000 *g*, 4 °C, for 30 minutes. Clarified lysates were supplemented with 5 mM MgCl2 and 4 mM imidazole, then applied to Ni-NTA resin (Qiagen) for batch binding, ∼1 hr at 4 °C. Following binding and flow-through, the resin was washed with 10 column volumes (cv) lysis buffer + 4 mM imidazole, 2 x 5 cv lysis buffer + 500 mM KCl + 1 mM ATP + 10 mM MgCl2 + 4 mM imidazole, then 2 x 5 cv M1 buffer (300mM NaCl, 25mM HEPES-NaOH pH 7.5, 2mM CaCl2, 10% glycerol) + 8 mM imidazole. Protein was eluted with M1 buffer + 300 mM imidazole, applied directly to equilibrated M1 α-FLAG nanobody resin (kind gift from Aashish Manglik lab), and batch bound ∼1 hr at 4 °C. The M1 resin was washed extensively with M1 buffer, and bound FLAG-HSF1-his6 proteins eluted 7 x 1cv with M1 elution buffer (150 mM NaCl, 10% glycerol, 20 mM HEPES pH 7.5, 2 mM EDTA, 0.1 mg/mL FLAG peptide). This eluate was concentrated and separated by size exclusion in SEC buffer (150 mM KCl, 10% glycerol, 25 mM HEPES pH 7.5, 0.5 mM Tris(3-hydroxypropyl)phosphine) (THP)) using a Superdex 200 10-300GL column (Cytiva). HSF1 monomer and trimer peaks were pooled separately, concentrated, and flash frozen.

HSF1 variants, HSF1(1-417), HSF1(cf), HSF1(1-417)-GFP, HSF1(1-417, cf)GFP, and single cysteine variants such as HSF1(cf, L125C, 1-417)GFP and HSF1(cf, L125C) were constructed, tagged, and purified as described above. This work uses cysteine-free SGFP283 in all cases where GFP is mentioned. This was achieved by introducing six point mutations to EGFP: C49S, C71M, M154T, V164T, S176G, and A207K. To fluorescently tag single cysteine HSF1 variants, M1 eluates were concentrated to 10-50 µM and supplemented with 100 µM Alexa Fluor 546 C5 maleimide (Invitrogen #A10258) or Alexa Fluor 647 C2 maleimide (Invitrogen # A20347) and rocked at 4 °C overnight. Monomers, trimers, and free dye were separated by SEC as described above.

In all instances in this work, the glucocorticoid receptor (GR) construct used in biochemical experiments was MBP-tagged GR ligand binding domain (MBP-GRLBD), corresponding to residues 521-777 with a stabilizing F602S mutation. MBP-GRLBD was expressed and purified as described in Kirschke et al., 201463, with slight modifications. MBP-GRLBD was present in both the flow through and an eluted peak from the anion exchange column, with equal purity. These were separately taken through the rest of the purification; the MonoQ-binding portion notably contained dimeric GR. Following, the final dialysis step, GR monomer fractions were re-run through a Superdex 200 16/60 HiLoad (Cytiva) column using 150mM KCl, 10% glycerol, 20mM HEPES pH 7.5, 0.5 mM TCEP, then concentrated and stored in that buffer.

Fluorescently labeled GR-Alexa488 was prepared by mixing 70 µM MBP-GRLB in its storage buffer with equal-molar Alexa Fluor™ 488 Carboxylic Acid, 2,3,5,6-Tetrafluorophenyl Ester (ThermoFisher #A30005) and rocking overnight at 4 °C. Protein and free dye were separated using an Superdex 200 10/300 GL column, and the protein peak collected, aliquoted, and flash-frozen. Labelling efficiency, calculated by comparison of sample absorbance on a NanoDrop (Thermo Scientific) at 280 nm and 495 nm, was ∼ 50%.

### Purification of chaperones and co-chaperones

Full-length, wild-type human Hsp90AA, Hsp90AB, HOP, p23, FKBP52, FKBP51, and Bag-1 bag domain (222-345), and yeast Ydj1, were expressed and purified as described previously44,50,63,^67^. HOP(ΔDP2) and Hsp90(D93N) were expressed and purified in the same manner as the full-length constructs. All proteins were stored in 150mM KCl, 10% glycerol, 25 mM HEPES pH 7.5, 0.5 mM THP. Hsp70 (*HSPA1A*), Hsp70(T204A), Hsp70(ΔEEVD), Hsc70 (*HSPA8*), DnaJB1, DnaJB1 variants, DnaJA1, NudC, Sugt1A, and Sugt1B were expressed and purified similarly to other (co-)chaperones with slight modifications. Genes encoding these constructs were cloned into pQiq vectors containing N-terminal His10-SUMO tags, transformed into BL21(DE3) or BL21(DE3) RIL *E. coli* and grown at 37 °C in terrific broth containing 50 µg/mL kanamycin sulfate (and + 35 µg/mL chloramphenicol for BL21(DE3) RIL cells). At reaching OD600 ∼ 1.0, cultures were cooled to 18 °C, induced with 200 µM IPTG, and expressed overnight. The next day, cells were harvested, resuspended in lysis buffer, and supplemented with 10 µg/mL lysozyme (Sigma Aldrich), 1.0 µg/mL bovine DNaseI, 1.0 mM phenylmethylsulfonate fluoride (PMSF, Thermo Fischer), cOmplete protease inhibitor (Roche), 3 mM β-mercaptoethanol (βMe), and 1 mM ethylenediaminetetraacetic acid-NaOH pH 8.0 (EDTA). Cells were homogenized on ice with a dounce, then lysed by 3x passage through an EmilsiFlex-C3 (Avestin) at 15,000 psi. Lysates were clarified by ultracentrifugation, and the supernatant supplemented with 1mM MgCl2 and 10mM imidazole. Proteins were eluted with 300 mM imidazole in ion exchange buffer A, containing 50 mM KCl, 5% glycerol, 0.5 mM THP, and either 30 mM Tris-HCl pH 8.5 or 30 mM MES-NaOH pH 6.0 (DnaJB1 constructs). Eluted proteins were digested with Ulp1cat while dialyzing overnight against Buffer A at 4 °C. Following digestion, DnaJB1 constructs were loaded onto a MonoS 10/100 GL anion exchange column (Cytiva), and eluted with a linear gradient of 50-500 mM KCl. The other proteins were loaded onto a MonoQ 10/100 GL anion exchange column (Cytiva), and likewise eluted with a linear gradient of 50-500 mM KCl. For all proteins, the final purification step was SEC using either Superdex 200 or Superdex 75 columns (Cytiva) in 150mM KCl, 10% glycerol, 25 mM HEPES pH 7.5, 0.5 mM THP.

### Purification of pBPA variants

HOP(E509pBPA)-His6, Hsp90AA(XpBPA) and Hsp90AB(XpBPA) variants were expressed and purified as the wild-type proteins, with slight modifications. The vectors pET151-HOP(E509pBPA)-His6, pET151-His8-V5-TEV-Hsp90AA(XpBPA) or pET151-His6-V5-TEV-Hsp90ABco(XpBPA) were co-transformed with pEVOL-pBpF43 into BL21(DE3) *E. coli* and grown at 37 °C in terrific broth + 50 µg/mL carbenicillin + 50 µg/mL chloramphenicol to OD600 ∼ 1.0-1.5, then supplemented with 0.7 mM *para*-benzoyl-phyenalanine (pBPA) and 0.02% w/v arabinose and cooled to 24 °C, then induced with 300 µM IPTG. Subsequent cell harvesting, lysis, Ni-NTA affinity purification, TEV digestion and dialysis, anion exchange chromatography, and SEC were performed as previously described, with the final buffer containing 150mM KCl, 10% glycerol, 25mM HEPES pH 7.5, and 0.5 mM THP.

### DNA binding and fluorescence polarization assays

Trimeric HSF1 was generated by heat shocking 10 µM purified HSF1 monomers at 41 °C for 15 minutes. Although HSF1 purified as trimers was also functional in these experiments, degradation products co-purified within the trimers, making measurements more variable between replicates.

HSE-containing DNA oligonucleotides with a 5’ fluorescein amidite modification (FAM-HSE) were used in fluorescence polarization (FP) assays. The labeled oligonucleotides were annealed to complementary DNA by heating to 95 °C then cooling 1 °C per minute from 80 °C to 25 °C. Oligos were typically stored in 150 mM KCl, 10% glycerol, 25mM HEPES pH 7.5, and 0.5 mM THP; similar results were achieved with NaCl in the buffer. FP experiments were performed in the same buffer with 5 nM FAM-HSE and 5 µM BSA. Trimeric HSF1, wt or variants, were mixed with equal volumes of buffer with 10 nM FAM-HSE and 10 µM BSA, then serially diluted at room temperature. FP was measured using either a Spectromax M5 plate reader (Molecular Devices) or a CLARIOstar plate reader (BMG Labtech). After equilibration, milli-polarization values were read for all concentrations and FAM-HSE without HSF1. Three or more independent experiments were conducted for each construct, and Kd values were calculated using GraphPad Prism using the single-site saturation binding function.

To monitor DnaBJ1 and Hsp70 activity by FP, 0.8 µM trimeric HSF1 (2.4 µM monomer equivalent), 100 nM 5’FAM-HSE, 5 µM DnaJB1 (monomer concentration), and 10 µM Hsp70 were co-incubated at 28 °C in a Spectromax M5 plate reader, and FP measure every 2 minutes. Reactions without HSF1 were used to show the baseline. To start reactions, 3 mM ATP + 4 mM MgCl2, or just mM MgCl2 were added, rapidly mixed, and returned to the plate reader.

### General procedures for FRET Measurements

|  |  |  |  |
| --- | --- | --- | --- |
| **Auto-inhibition:**<br>monomer – high<br>trimer – low | Ex: 460 nm (GFP) | **Oligomerization:**<br>monomer – low<br>trimer – high | Ex: 540 nm (Alexa 546) |
|  | Em: 570 nm (Alexa 546) |  | Em: 670 nm (Alexa 647) |
|  | Ex: 460 nm (GFP)<br>Em: 520 nm (GFP) |  | Ex: 540 nm (Alexa 546)<br>Em: 610 nm (Alexa 546) |

To determine optimal wavelengths for monitoring HSF1 conformational changes and oligomerization, excitation and emission scanning were performed with HSF1(fret) monomers and trimers (Figure S1B). Measurements were taken using Spectromax M5 or CLARIOstar plate readers, with samples in clear-bottom, black, 96-well or 384-well plates (Corning) sealed with HD Clear packing tape (Duck). For subsequent measurements, wavelengths were chosen to maximize the ratio of monomer signal to trimer signal. Thus, HSF1 auto-inhibition FRET signal was calculated as (emission 570nm / emission 520nm) with excitation at 460 nm. HSF1 oligomerization signal was calculated as (emission 670nm / emission 610nm) with excitation at 540 nm. Notably, labeling efficiency of HSF1 single cysteine variants was variable, making values not directly comparable between some experiments. To calculate averages from biological replicates, data were scaled to the range between the trimer and monomer values for each batch of HSF1(fret).

### Reconstitution of HSF1(fret) activation and inactivation

For measurements of HSF1 trimerization rates (Figures S1, S3), concentrated HSF1(fret) monomers at room temperature were diluted into pre-warmed buffer in pre-warmed plates at the indicated temperature, plates sealed, and measurements begun immediately. Initial rate calculations were calculated from scaled data (see above) in the 5-to-15-minute post-dilution window. For subsequent experiments using HSF1(fret) trimers, 10 µM HSF1 monomers, comprised of equimolar HSF1(cf,1-417, L125C-Alexa Fluor 546)-GFP and HSF1(cf, L125C-Alexa Fluor 647), were heat 15 minutes at 41 °C in a hot water bath with rotation at 300 rpm. Extent of trimerization was confirmed by size exclusion chromatography.

To monitor chaperone-driven changes to HSF1 state, HSF1(fret) monomers or trimers, stored on ice, were mixed with other relevant components at room temperature, transferred to plates, sealed, and equilibrated in plate readers while initial measurements were taken (e.g., Figure 1c,d). Measurements were carried out at 28 °C unless otherwise noted, in buffer containing 150mM KCl, 4 mM MgCl2, 10% glycerol, 25mM HEPES pH 7.5, and 0.5 mM THP. To start reactions, ATP was spiked into 3 mM final concentration and mixed rapidly.

### UV photo-crosslinking in HSF1-chaperone reconstitutions

Components, on ice, were assembled into reactions at room temperature in UV-transparent PCR tubes (Thomas Scientific #1149Z79), using protein storage buffer (150mM KCl, 10% glycerol, 25mM HEPES pH 7.5, and 0.5 mM THP) as the base buffer. To start reactions, ATP was added to 3 mM and MgCl2 to 4 mM, unless otherwise noted, and the PCR tubes were incubated in a thermocycler. Final concentrations of components are noted in figure legends. In experiments with inhibitors, stock solutions and controls were made to ensure all samples had equal final concentrations of DMSO. If components were added to some samples during reactions, buffer was added to the other samples. Reactions were typically carried out at 28 °C, deviations are noted in figure legends. When the desired endpoint was reached, reactions were moved to ice for 5 minutes, then photo-crosslinked under a UV lamp (Alpha Thera) at 4 °C for 1 hr.

In most cases, crosslinked reactions were directly mixed 3+1 with 4x NuPAGE LDS sample buffer (Invitrogen # NP0007) containing 4% v/v β-Me and heated 10 minutes at 80 °C prior to PAGE analysis. For in-gel fluorescence imaging, gels were imaged with a Typhoon FLA 9000 (GE Healthcare) or GEL-DOC XR+ (Bio-Rad), then stained with colloidal Coomassie blue and destained with water for total protein imaging on an Azure 600 imaging system. For western blotting, samples were transferred to PVDF using an iBlotTM 3 transfer system, blocked with PBS-TWEEN20 with 3% w/v BSA. FLAG-HSF1 was imaged using anti-FLAG-HRP (Sigma #A8592), and MBP-GRLBD was imaged using anti-maltose binding protein-peroxidase (Sigma #A4213).

### Sample preparation for cryo-EM

Loading state complexes were prepared by reconstitution of 8 µM FLAG-HSF1(wt) monomers, 30 µM Hsp90(D93N), 8 µM DnaJB1, 30 µM Hsp70, and 15 µM HOP(E509pBPA) at 28 °C for 80 minutes followed by photo-crosslinking for 1 hr at 4 °C. Total reactions, 40 µL, were then directly injected on to a Superose 6 3.2/300 size exclusion column (Cytiva) on an Ettan LC system (Amersham) at 4 °C in loading state buffer (75 mM KCl, 30 mM HEPES-NaOH pH 7.5, 0.5 mM THP, 2% glycerol, 0.2 mM ADP, 0.05% beta-OG) (Figure S12). Peak fractions, previously analyzed by α-FLAG western blotting total protein staining, were directly transferred to ice and immediately used for grid preparation. HSF1 concentration in the samples was estimated at 0.5 – 1.0 µM from parallel experiments using HSF1(cf, Y225C-alexa647).

Maturation state complexes containing HSF1 were prepared by reconstitution of 8 µM twinStrep2-HSF1(wt) monomers, 30 µM Hsp90(W320pBPA or E451pBPA), 18 µM DnaJB1, 36 µM Hsp70, and 18 µM HOP(wt) at 28 °C for 40 minutes, addition of 30 µM p23 and 10 mM NaMo4 and incubation for an additional 40 minutes. Samples were moved to ice, then photo-crosslinked for 1 hr at 4 °C. Next, 50 µL of crosslinked samples were applied to 50 µL of strep-tactin-XT superflow high-capacity resin (IBA #2-4030-025) in capped 1mL gravity flow columns on ice. Resin was pre-equilibrated with Strep wash buffer, 150 mM KCl, 25 mM HEPES-NaOH pH 7.5, 0.5 mM THP + 0.01% DDM, 10 mM NaMoO4, 2 mM MgCl2, 2 mM ATP supplemented with 60 µM Bag-1 bag domain to promote Hsp70 dissociation. Protein was incubated with the resin for 1 hr, and resuspended every 10 minutes by gentle shaking. Then the resin was washed 8 x 1cv with ice cold wash buffer, and bound proteins eluted with 8 x 1cv elution buffer (75 mM KCl, 30 mM HEPES-NaOH pH 7.5, 0.5 mM THP, 50 mM biotin-NaOH, 0.005% DDM, 2 mM ATP, 2 mM NaMoO4, 2 mM MgCl2). Strep eluates were concentrated using Vivaspin 50kDa MWCO concentrators (Cytiva #28932236) to final HSF1 concentration 0.5 – 1.0 µM, estimated from parallel experiments using twinStrep-HSF1(cf, 1-417, Y225C-alexa546)-GFP.

### Cryo-EM data collection, processing, and model building

Microscope parameters and data collection parameters were the same for all data sets. Micrographs were collected using SerialEM with a Titan Krios (Thermo Fischer Scientific equipped with a K3 Summit direct electron detector (GATAN) at 105,000X magnification. The pixel size was 0.8189 Å with a dose rate of 16e-/pix/s, 2 s exposure time and 0.025 s frame rate. Micrographs were motion corrected using UCSF Motioncor2 (Zheng, 2017)^84^ and CTFs were initially calculated using CTFFIND4.185. Subsequent processing was carried out in cryoSPARC^86^. Micrographs were curated by CTF cutoff and ice thickness.

For the Hsp90(E451pBPA)2-p23-HSF1 maturation complex, particles were picked using templates generated from a previous structure of Hsp902-p23 (EMD-23006)^44^. Approximately 4.5 million picked particles were triaged by two rounds of 2D classification, yielding ∼1.9 million particles which appeared to contain protein. One tenth of these particles were used for *ab initio* reconstruction (3 classes), yielding one class clearly containing a closed Hsp90 dimer and two classes with unclear structure. All particles accepted by 2D classification were sorted into these classes by heterogeneous refinement, and the ∼ 1.0 million particles in the good class were re-extracted and aligned by non-uniform refinement, with a nominal resolution of 2.5 Å. In this initial consensus map the closed Hsp90 dimer was well-resolved, however, density for p23 was sparse and density in the lumen of Hsp90 was difficult to interpret.

Focused 3D-classification on the Hsp90 NTDs and p23 resolved a class (∼333k particles, ∼2.7 Å resolution) with one copy of p23 in which the map was also more interpretable in the Hsp90 lumen. Further attempts to classify on the Hsp90 MDs and client HSF1 revealed that other conformations were present but decreased map interpretability. While it is possible that this also reflects Hsp90(E451pBPA) crosslinking to p23, those species were a very minor fraction of the final sample frozen (Figure S12A) whereas Hsp90-Hsp90 and Hsp90-Hsp90-HSF1 species were enriched. Although the sample used wild-type Hsp90 with ATP and NaMoO4, electron density in the NTDs more strongly resembles ATP than ADP-MoO4, likely reflecting partial ATP hydrolysis.

For the Hsp90(E451pBPA)2-HSF1(1-417)-GFP complex, particles were identified by Gaussian blob picking. The initial stack of ∼2.9 million particles was cleaned by one round of 2D classification; all particles in classes with distinguishable protein were kept (∼909 k particles). A random subset of these particles were used for *ab initio* reconstruction with varying numbers of classes. In all cases the best class contained a closed Hsp90 dimer; some such classes also appeared to contain p23 and additional density near the Hsp90 MDs. The classes were combined and refined to an initial consensus map (438 k particles, ∼3.3 Å nominal resolution) comprising a closed Hsp90 dimer, p23 and additional barrel shaped density that roughly accommodated a GFP molecule. Focused classification on the Hsp90 MDs and GFP suggested that the GFP was relatively mobile, however, one class appeared to correspond to a favored conformation (72 k particles) in which p23 was absent and the GFP barrel was perpendicular to the Hsp90 dimer axis. Attempts to resolve the linker between the GFP and the HSF1 bound in the Hsp90 lumen were unsuccessful. The refined class with GFP had a nominal resolution of ∼3.7 Å, and while the local resolution of the GFP was much lower, no other component in the sample reconstitution made sense to fit in the map.

To pick particles of the Hsp90(D93N)2-Hsp702-HOP(E509pBPA)-HSF1 loading complex, templates were generated from the map of the Hsp90-GR loading complex (EMD-23050)^45^. In template generation, the map was thresholded to exclude density for client GR, binarized, then low-pass filtered to 15 Å. Picking with 50 templates yielded ∼2410 k extracted particles, and most particles (∼1741 k) were kept following one round of 2D classification. One tenth of these particles were used for *ab initio* reconstruction (6 classes), yielding one class clearly containing an Hsp90 loading complex and five classes with unclear structure. All particles accepted by 2D classification were sorted into these classes by heterogeneous refinement, and the ∼ 777 k particles (45%) in the best class were re-extracted and aligned by non-uniform refinement, with a nominal resolution of ∼3.2 Å.

A second, symmetric, copy of HOP was partially present and not (wholly) an artifact of mis-alignment. These particles were removed by 2 rounds of heterogeneous refinement between two classes with one or two copies of HOP. The nominal resolution of the cleaned consensus map was still ∼3.2 Å (395 k particles). Varying 3D classification strategies yielded classes missing one copy of Hsp70, without one or both Hsp90 NTDs, or with all components previously observed. Notably, variable Hsp70 SBD orientations were observed as well, and further efforts made on the subset of particles (129k) in which a full copy of one Hsp70 SBD was resolved in a complete loading complex.

Focused classification and local refinement were used to improve areas of interest in the loading state consensus map. In particular, two overlapping regions with client (HSF1) interaction sites were improved by first recentering and re-extracting particles in the full-SBD stack (129k) then performing particle subtraction and local refinement. Composite maps were made using the Phenix^87^ tool Combine Focus Maps following initial model docking (below) into the consensus map. The composite map was used for later stages of model refinement because it is the map used for interpretation of the model and other results.

Model building and refinement were carried out in UCSF ChimeraX^88^, Coot^89^, and Phenix^87^. Initial docking of structures into maps was performed in ChimeraX using Hsp90 and p23 from PBD #7KRJ^44^ for closed states, and Hsp90, Hsp70 NBDs, and HOP from #7KW7^45^ for the loading state. EGFP structure #6YLQ^90^ was used as the starting point for HSF1(1-417)-GFP, and appropriate mutations were modeled in Coot. For the Hsp70 SBD seen in the loading state the initial model docked was the crystal structure of the human Hsp70 SBD bound to a substrate peptide, #4PO2^91^. At residues where pBPA substitutions were made, the crosslinked adducts are visible in the maps, especially at Hsp90(E451pBPA) in the in model building. Following initial docking, alternating rounds of manual adjustment in Coot and automated refinement using Phenix Real-space refinement were used to improve the model. Reference model restraints were used for low-resolution regions, and secondary structure restraints used throughout. Sharpened maps from cryoSPARC and Phenix Autosharpen map were used to aid model building.

### Crosslinking mass spectroscopy Sample preparation

Loading and maturations state complexes containing Hsp90-HSF1 or HOP-HSF1 adducts crosslinked with pBPA were made as described above under cryo-EM sample preparation. For loading state samples, complexes were prepared using both Hsp90(E451pBPA) and Hsp90(D93N, E451pBPA). For Maturation state complexes, Hsp90(E451pBPA) was used. For HOP-HSF1 crosslinking, HOP(E509pBPA) was used with Hsp90(wt) or Hsp90(D93N). Additionally, samples were prepared using HSF1(wt), DnaJB1, Hsp70, and HOP(E509pBPA) without Hsp90; this was done analogously to the loading state samples.

Separately, HSF1-Hsp40-Hsp70 complexes were prepared (Figures 4 and S4A,B). Trimerized HSF1(1-417) (6 µM monomer equivalent) was incubated with 15 µM DnaJB1 and 15 µM Hsp70 in buffer containing 150 mM KCl, 25 mM HEPES pH 7.5, 10% glycerol, 0.5 mM THP, and 5 mM MgCl2 for 1 hr. Then ATPγS was added to 5 mM and the incubation continued 4 hrs longer. Lastly, DSSO was added to 2 mM (100 mM stock in DMSO), and the incubation continued 1 hr longer. The reaction was quenched with 20 mM Tris-HCl pH 7.5 and run directly on a Superose 6 3.2/300 GL column in 50 mM KCl, 50 mM NaCl, 20 mM HEPES-NaOH pH 7.5, 0.5 mM TCEP, 1% glycerol. Peak fractions were analyzed by SDS-PAGE and those containing high molecular weight species were concentrated for further use.

### Liquid Chromatography

Liquid chromatography was performed using a Vanquish Neo UHPLC system (Thermo Fisher Scientific) configured in a direct injection format. Peptides were separated on the Vanquish Neo with an Aurora Ultimate C18 120 Å, 1.7 µm, 75 µm x 60 cm UHPLC column (Ion Opticks). During LC separations, mobile phase A (MPA) was 0.2% FA in water and MPB was 80% ACN in water with 0.2% FA. For profiling of the proteome, a 90-min gradient ramped, at a flow rate of 300 nL/min, from 0-11% MPB from 0 – 2 min, 11-45% MPB from 2 – 68 min, 45 – 58% MPB from 68 – 78 min, 58 – 99% MPB from 78 – 80 min, and held at 99% MPB to 90 min before the column was washed and re-equilibrated at 0% MPB for four column volumes. During peptides separations, the LC column was held at 50 °C.

### Mass Spectrometry

Data-dependent acquisition (DDA) was performed on a Thermo ScientificTM Orbitrap EclipseTM TribridTM mass spectrometer system (Thermo Fisher Scientific, San Jose, USA). Precursors were ionized using electrospray ionization at 2 kV with respect to ground. The inlet capillary was held at 275 °C, and the ion funnel RF was held at 30%. During DDA experiments, all MS1 survey scans were acquired at a resolving power of 60,000 in the Orbitrap analyzer with a scan range of *m/z* 380 – 2000, maximum injection time of 50 ms, and AGC target of 1,000,000 charges.

Monoisotopic precursor selection was enabled for peptide isotopic distributions. Dynamic exclusion was set to exclude resampling of precursors ±10 ppm within 30 s. MS2 scans were conducted on precursors of a charge state z = 2-8 for a 2 s cycle time. Precursors were isolated with an isolation width of 0.9 Th using the quadrupole. MS2 scans were conducted in the Orbitrap at a resolving power of 60,000 at 200 m/z and an AGC target of 100,000 with a maximum injection time of 118 ms. Stepped HCD normalized collision energy (NCE) was set to 20%, 30%, and 35%. Scan range was set to Auto.

## Data Analysis

DDA .raw files were converted to .mzML files using MSConvert^92^. Data were further processed using MeroX^93^. To identify crosslinked species, data analysis was performed similarly as described in Kolhe et al., 202331. Settings denote proteolytic cleavage C-terminal to lysine and arginine not before proline with up to 2 missed cleavages. A FASTA file containing the sequences for DnaJB1, Hsp70, Hsp90, p23, and HSF1 was downloaded from UniProt on April 9, 2024. Peptide length of 7 to 52 amino acids was specified. Alkylation of cysteines and site-specific BPA incorporation were fixed while oxidation of methionine and protein N-terminal acetylation were variable modifications. Precursors, peptide identifications, and proteins were filtered to maintain 1% FDR.

## Assessing conformational state of HSF1(fret) in HspG0 complexes

HSF1(fret) monomers and trimers were prepared as described above. Chaperone complexes were prepared using 6 µM HSF1(fret) monomers, 30 µM Hsp90(W320pBPA), 24 µM HSp70, 6 µM DnaJB1, and 15 µM HOP. Loading state complex assembly used HOP(E509pPBA), whereas maturation state structure. Nonetheless, the maps also do not reflect the geometry of a single crosslink at near 100% completion, and so the monomer PBF was used as the residue maturation state assembly used HOP(wt), 30 µM p23, 10 mM NaMoO4. Components were incubated 80 minutes at 28°C, with p23 and NaMoO4 added after 40 minutes to maturation complexes. Next, reactions were moved to ice for 5 minutes and exposed to UV for 1 hr at 4 °C. Crosslinked samples were directly injected onto a Superose 6 3.2/300 SEC column in buffer (75 mM KCl, 30 mM HEPES-NaOH pH 7.5, 0.5 mM THP, 2 mM MgCl2) for maturation complexes, or buffer plus 0.2 mM ADP for loading complexes. Retention volume of HSF1(fret) was monitored by absorbance at 640 nm and 560 nm, while total protein was monitored at 280 nm. Fractions (50 µL) were collected in 96 well plates and HSF1(fret) fluorescence measured in a CLARIOstar plate reader.

## HSF1 Bait-Prey pulldowns

Prey FLAG-HSF1, 3 µM, was reconstituted with 20 µM Hsp90, 15 µM Hsp70, 4 µM DnaJB1, 10 µM HOP, and where indicated, 20 µM p23. Variants of these components are indicated in figure legends. Chaperone components excluding p23 were assembled at room temperature with 3 mM ATP, and reactions started with the addition of HSF1 and transferred to 28 °C for 40 minutes. Then, where indicated, p23 and 10 mM NaMoO4 were added, and all reactions incubated at 28 °C an additional 40 minutes. Reactions were moved to ice for five minutes, then exposed to UV for 1 hr at 4 °C. Following crosslinking, twinStrep-HSF1 was spiked into all reactions to 3 µM, and samples then heated at indicated temperatures (25 °C to 42 °C) for 20 minutes. Reactions were moved to ice and quenched by addition of 5 mM EDTA, then applied to Strep-Tactin resin on ice.

Pulldowns were performed similarly to those described above for maturation state complexes. Initial experiments were performed with twinStrep-HSF1(wt) as bait and FLAG-HSF1(wt) as prey, and HSF1 pull-down (trimerization) efficiency was judged western blotting with anti-FLAG-HRP and Strep-Tactin-HRP (IBA #2-1502-001). Subsequently, FLAG-HSF1(cf, Y225C-Alexa647) was used as prey and twinStrep-HSF1(cf, 1-417, Y225C-Alexa546)-GFP was used as bait. For experiments with HSF1 truncations, FLAG-tagged and otherwise wild-type constructs were used as baits.

## DNA binding by chaperone-bound HSF1

For DNA binding of HSF1 in maturation state complexes (Figures 6A,B) the HSF1 monomers used consisted of a mixture of twinStrep-HSF1(wt) and FLAG-HSF1(cf, Y225C-Alexa546). For controls with just HSF1 trimers, 8 µM monomer mixture was heated 15 minutes at 41 °C, 300 rpm to induce trimerization. HSF1-Hsp90 maturation complexes were assembled with 24 µM Hsp90(W320pBPA), 20 µM Hsp70, 4 µM DnaJB1, 12 µM HOP, and 24 µM p23. Otherwise, maturation complex formation and crosslinking were carried out as described for bait-prey experiments, above.

For Figure 6a, following crosslinking, samples were heat-shocked for 20 minutes, 41 °C, 300 rpm, then returned to ice. As indicated, fam-HSE or fam-HSE90AA with or without 40 µM Bag-1 bag domain were added and incubated on ice for 5 minutes. Samples were then directly injected onto a Superose 6 3.2/300 column in 75 mM KCl, 30 mM HEPES-NaOH pH 7.5, 0.5 mM TCEP, 0.1 mM ADP, 0.01% DDM. Elution fractions were directly scanned for HSF1 (Alexa Fluor 546) and DNA (fitc) fluorescence in a CLARIOstar plate reader.

For Figure 6b, samples were assembled, crosslinked, and heat shocked as described above. Bag-1 bag domain (40 µM) was added to all samples on ice, and samples were applied to Strep-Tactin resin as described for bait-prey experiments and cryo-EM sample preparation, with small changes. Here, wash and elution buffers contained 150 mM KCl, 25 mM HEPES-NaOH pH 7.5, 10% glycerol, 0.5 mM THP, 0.01 % DDM, 1mM EDTA, and 2 mM ATP, with 50 mM biotin-NaOH in the elution buffer. Strep-Tactin eluates were concentrated then incubated with 2.0 µM fam-HSE or 1.0 µM fam-HSE90AA on ice prior to SEC and analysis as described above.

For figure 6C, loading and maturation state complexes were treated as described above for figure 6A, with small adjustments. Samples that were not heat shocked were kept on ice for the corresponding incubation time. For the loading state samples, 24 µM Hsp90(D93, E451pBPA), 20 µM Hsp70, 4 µM DnaJB1, and 12 µM HOP(E509pBPA) were used. SEC elution fractions were analyzed by SDS-PAGE, in-gel fluorescence, and total protein staining.

## Notes

### Competing Interest Statement

The authors have declared no competing interest.

